# Spatial dynamics and vaccine-induced fitness changes of *Bordetella pertussis*

**DOI:** 10.1101/2021.11.01.466439

**Authors:** Noémie Lefrancq, Valérie Bouchez, Nadia Fernandes, Alex-Mikael Barkoff, Thijs Bosch, Tine Dalby, Thomas Åkerlund, Katerina Fabianova, Didrik F. Vestrheim, Norman K. Fry, Juan José González-López, Karolina Gullsby, Adele Habington, Qiushui He, David Litt, Helena Martini, Denis Piérard, Paola Stefanelli, Marc Stegger, Jana Zavadilova, Nathalie Armatys, Annie Landier, Sophie Guillot, Samuel L. Hong, Philippe Lemey, Julian Parkhill, Julie Toubiana, Simon Cauchemez, Henrik Salje, Sylvain Brisse

## Abstract

Competitive interactions between pathogen strains drive infection risk. Vaccines are thought to perturb strain diversity through shifts in immune pressures, however, this has rarely been measured due to inadequate data and analytical tools. *Bordetella pertussis* (*B. pertussis*), responsible for 160,000 deaths annually^1^, provides a rare natural experiment as many countries have switched from whole cell vaccines to acellular vaccines, which have very different immunogenic properties^2,3^. Here we use 3,344 sequences from 23 countries and build phylogenetic models to reveal that *B. pertussis* has substantial diversity within communities, with the relative fitness of local genotypes changing in response to switches in vaccine policy. We demonstrate that the number of transmission chains circulating within subnational regions is strongly associated with host population size. It takes 5-10 years for individual lineages to be homogeneously distributed throughout Europe or the United States. Increased fitness of pertactin-deficient strains following implementation of acellular vaccines, but reduced fitness otherwise, can explain long-term genotype dynamics. These findings highlight the role of national vaccine policies in shifting local diversity of a pathogen that still poses a large burden on global public health.

## MAIN

The role of local population immunity in driving ecological interactions between strains from the same disease system remains a foundational question of infectious disease dynamics and ecology. However, immunity’s role in determining genetic diversity has rarely been explored as doing so requires long-term genetic data as well as perturbations in population immunity. We also need appropriate analytical tools that can quantify changes in lineage fitness. *B. pertussis* provides a rare natural experiment as many countries have switched from whole cell vaccines (WCV) to acellular vaccines (ACV), which have very different immunogenic properties^2,3^. *B. pertussis* is the causative agent of whooping cough, responsible for an estimated 160,700 deaths each year^1^. Despite long-standing immunization programs and high vaccine coverage levels, *B. pertussis* continues to circulate endemically, including with increased transmission in recent decades, the reasons for which remain unclear^4^. As with many bacterial systems, the study of *B. pertussis* is complicated by frequent asymptomatic carriage and long-term endemic co-circulation of multiple lineages^5–10^. Understanding the drivers of strain dynamics has important public health implications as different genotypes have been linked to differences in virulence, and with the duration and efficacy of vaccine protection^11–13^.

We established a consortium of national reference laboratories to sequence the whole genome of 1,331 isolates from 12 European countries. With publicly available genomes, this generated a dataset of 3,344 genomes from 23 countries and 5 continents covering an 85-year period (Figure 1A, Table S1-2). Subnational location was recorded for 97% of genomes. We characterised the diversity of *B. pertussis* across different spatial scales (within-district, within-country, intra-continental and global), estimated its rate of expansion, and assessed the importance of the local population size in driving the diversity of circulating lineages. We separately developed an analytical framework that estimated the relative fitness of different lineages, and characterised fitness changes following switches in the local vaccine being used. Our analytical approach allowed us to answer critical questions linked to *B. pertussis* diversity within and between locations, long-term genotype dynamics and a role for national vaccine policy in driving genotype changes.

**Figure 1:**
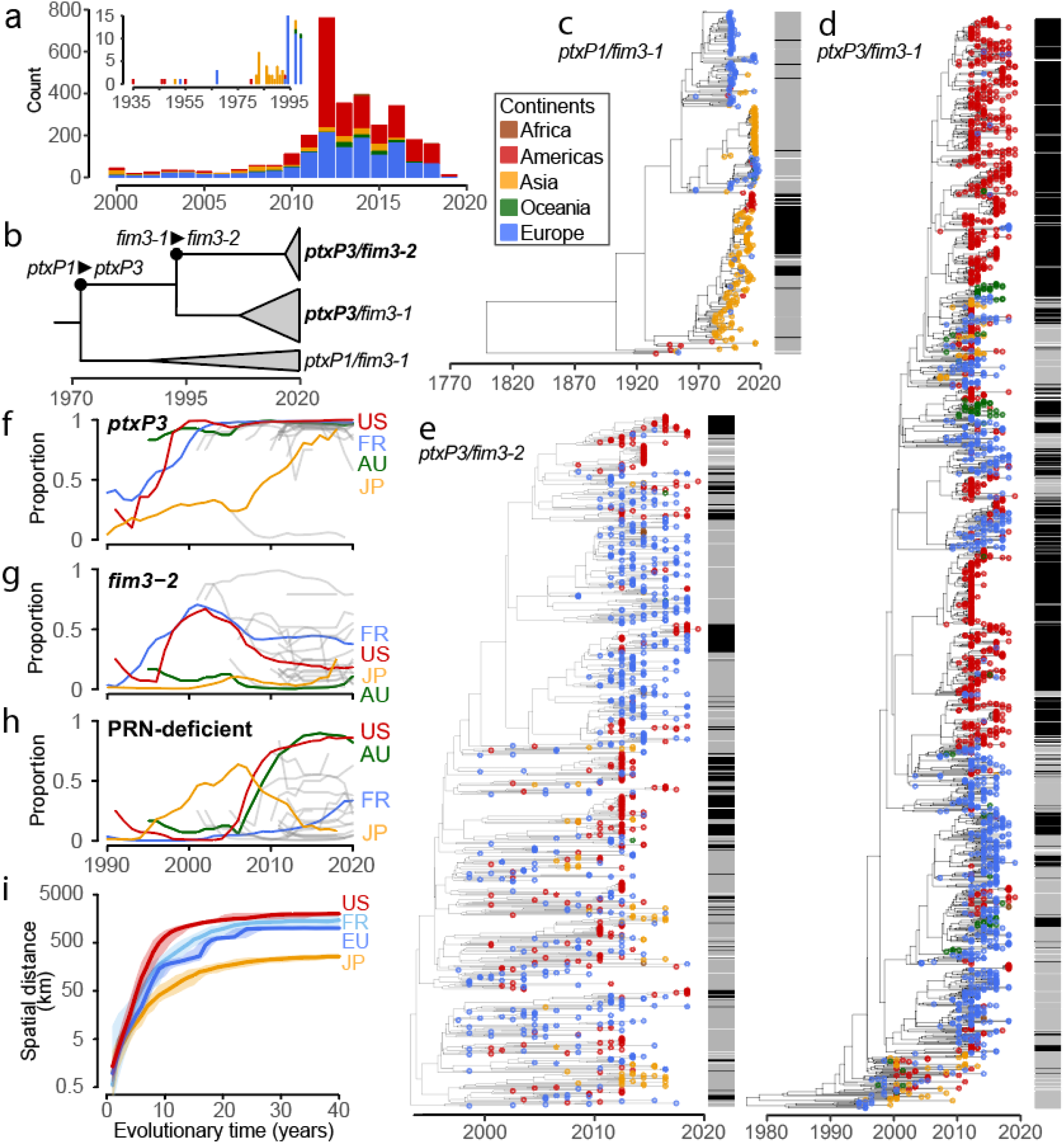
Origin of sequences and genetic diversity of *Bordetella pertussis*. (A) Number of sequences by continent, as a function of time (see Table S1 for further details). (B) Schematic of the evolutionary relationship of the *ptxP* and *fim3* alleles. (C-E) Maximum clade credibility trees for the different genotypes: *ptxP1/fim3-1* (C), *ptxP3/fim3-1* (D) and *ptxP3/fim3-2* (E). Branch tips are colored by the continent of collection. Horizontal bars denote pertactin (PRN) expression (black: deficient expression, grey: wild type expression, white not known) (F-H) The full tree is available on the MicroReact platform [URL to be provided on acceptance]. Temporal trends in strain frequencies, computed on rolling 7-year windows, for the *ptxP3* (F) and *fim3-2* (G) alleles, and PRN-deficiency (H). (I) Median spatial distance between *B. pertussis* pairs from different locations (EU: Europe; FR: France; JP: Japan) separated by different evolutionary times. Global average over the first 5 years is presented in Figure S1. The shaded area represents 95% CIs.

### Temporal dynamics of *B. pertussis* genetic diversity

To characterise genetic diversity, we built a time-resolved (Figure 1B-E). In addition, we identified the genotype of each isolate based on the allele of the promoter region of pertussis toxin (PT, with promoter forms *ptxP1* or *ptxP3*) and type 3 fimbrial protein gene (*fim3-1* or *fim3-2*)^11,14^. PT is the major toxin produced by *B. pertussis* and is responsible for most of the systemic symptoms associated with pertussis disease^15^. Isolates with a *ptxP3*-type promoter have been suggested to produce more PT than those with a *ptxP1*-type promoter^11^. Fim3 is one of the two fimbriae produced by *B. pertussis* and is involved in adhesion to host cells^16^. A polymorphism of the *fim3* gene has been reported to have occurred mainly for isolates with a *ptxP3*-type promoter^17^. In addition to these genotypes, we also identified whether each isolate was capable of producing the immunogenic surface protein pertactin (PRN-positive or PRN-deficient), a key target of vaccine-induced immunity^18^. We found that in nearly all countries, the proportion of *ptxP3* isolates increased from 5-20% in the early 1990s to >80% by mid-2000s (Figure 1F). By contrast, *fim3* allele distribution has been more steady over the years (Figure 1G). Finally, PRN-deficient strains have displaced PRN-positive strains in most countries (Figure 1H). These findings are consistent with previous efforts that identified changing patterns of genotype diversity^17^.

### *B. pertussis* spread across spatial scales

To track the spatial spread of *B. pertussis*, we compared the evolutionary distance of pairs of sequences with their spatial distance. We found a strong log-linear relationship for up to five years, with consistent patterns observed in different countries (Figure 1I, Figure S1). Adjusting for uneven sampling by location and year, we estimated that pairs of isolates separated by a year of evolutionary time from within the same country were separated by an average of 2.1km (95%CI: 1.1-3.8), rising to 41.5km (95%CI: 29.2-60.7) when separated by 5 years (Figure S1), with consistent patterns obtained with alternative approaches for the definition of population centroids (Figure S2).

Recent findings from household transmission studies have identified frequent subclinical transmission of *B. pertussis*^*10,19*^. Consistent with such widespread subclinical transmission maintaining strain diversity, we found that despite strong overall patterns of spatial structure in sequences, the majority of sequence pairs within any location at any time were only distantly related. At both national and subnational levels, fewer than one in twenty sequence pairs coming from the same year had a Most Recent Common Ancestor (MRCA) within the prior two years (Figure 2A, Figure S3); with the vast majority of pairs separated by more than 20 years of evolutionary time.

**Figure 2:**
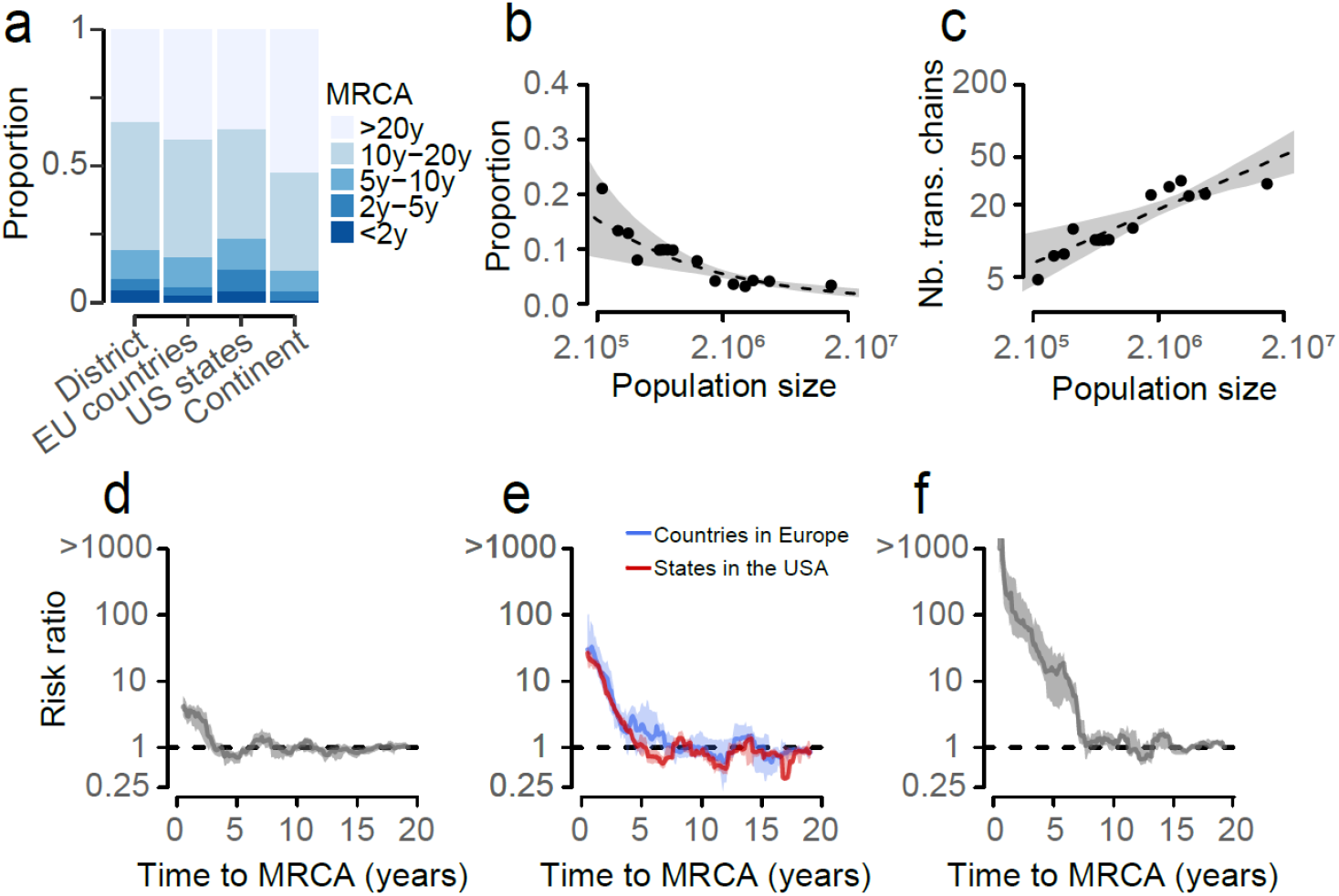
*Bordetella pertussis* diversity within and between locations. **(A)** Breakdown of the MRCA separating pairs of *B. pertussis* isolates from within the same year, for different locations: within districts (12 European countries, Iran and Japan), within European countries, within US states and within continents. **(B)** Proportion of pairs within a region that belong to the same transmission chain (defined as MRCA<2y), as a function of population size (average from 19 countries). Proportions are computed for rolling windows of population sizes. Dots represent the data and the dashed line represents model fit assuming an exponential relationship between the two and the grey shaded region 95% confidence intervals. **(C)** Number of transmission chains within regions, as a function of the population size. Numbers are computed for rolling windows of population sizes. Dots represent the estimates from the data and the dashed line represents model fit and the grey shaded region 95% confidence intervals. **(D-F)** Relative risk that a pair of bacteria have a MRCA within a defined period, when coming from the same versus different district in France (D), same versus different country in Europe (E, blue), the same versus different state in the US (E, red) or the same versus different continent (F). The shaded regions in (D-F) represent 95% confidence intervals.

To further explore lineage diversity at subnational levels, we considered pairs of individuals isolated within the same year and having an MRCA of under 2 years to be part of the same transmission chain. On average 4.3% of pairs (95%CI: 3.6-5.0) within each region (defined here as the smallest subnational administrative unit we have for each country; average population size: 6.5 million, average area: 110,000 km^2^) were from the same transmission chain (Figure 2B). There were an average of 23.2 discrete chains (95%CI: 20.0-28.0) circulating within a region. However, the number of chains depends strongly on population size (Figure 2C); each increase in log-population size was associated with a 0.45 (95%CI: 0.23-0.61) increase in the number of circulating chains, with consistent patterns across the different continents (Figure S4). Increases in population density were also associated with small increases in the number of circulating chains (Figure S5).

To characterise the longer-term spread of lineages, we calculated the probability that pairs from the same location were separated by an MRCA of specific time intervals, relative to pairs that come from different locations. In France, where we have the greatest spatial resolution, sequences from the same district were 2.6 times (95%CI: 1.5-3.9) as likely to have an MRCA within the two prior years as sequence pairs from different districts (Figure 2D). We found similar spatial clustering in other countries (Figure S6). Further, pairs of sequences coming from the same European country were 11.6 times (95%CI: 5.4-19.3) as likely to have an MRCA within the prior year than sequence-pairs coming from different European countries. We observed a similar level of spatial dependence between different US states as between European countries (Figure 2E). Overall, it takes approximately 2-3 years for a lineage to be well mixed within a European country (i.e., spatial structure is lost). Similarly, it takes *B. pertussis* 5-9 years to be well mixed across Europe, 4-6 years to be well mixed across the US and 8-12 years to be well mixed across different continents (Figure 2D-F, Figure S7). As we considered the relative probability of MRCA within and between locations for isolates sampled from the same year, our results are not affected by biased sampling between locations or in time^20^.

### Characterisation of the fitness of the different genotype, by vaccination era

In order to explain the observed changes in genotype in each location, we developed a time-series model that estimates the annual proportion of isolates that were of each genotype in each country. We use a single framework to fit the data from all the countries at the same time. We assumed that the relative fitness of the different genotypes were the same across countries but allowed for a change in fitness following the switch from WCV to ACV, which occurred at different times in different countries (Figure S8). This simple model was able to recover the time-series (Figure 3A-D, Figure S9), and the observed number, proportion, and annual relative change in genotypes across countries (Figure 3E-G).

**Figure 3:**
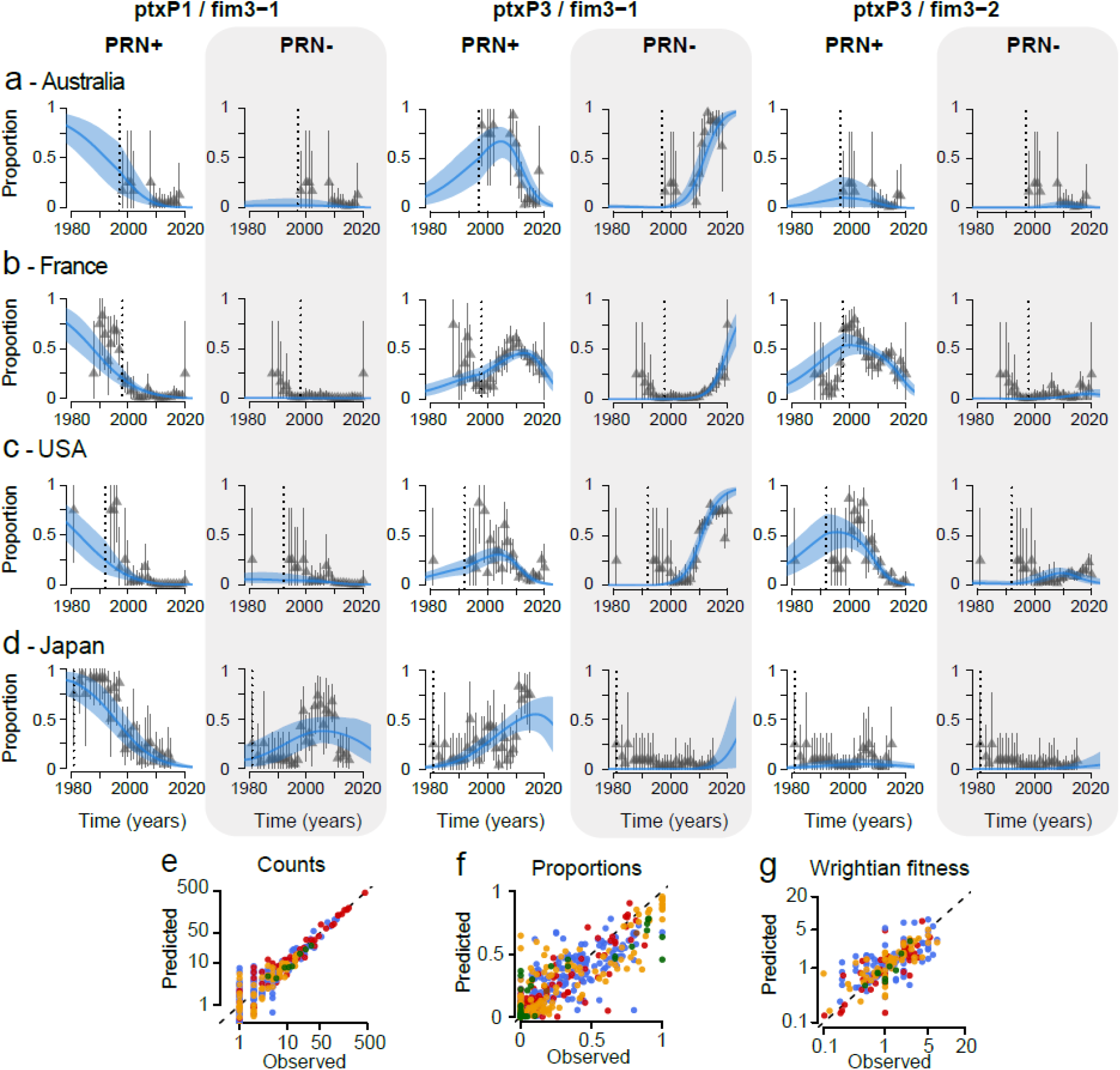
Changing genotype distributions of *Bordetella pertussis* by country. (A-D) Fits of the proportion of each genotype for 4 example countries: Australia (A), France (B), USA (C) and Japan (D) (other countries’ fits can be found in Figure S9). Grey triangles represent data, grey bars denote binomial 95% confidence intervals. Blue lines and shaded areas represent the median and 95% credible interval of the posterior. Vertical dotted line denotes the year of the first ACV introduction, for each country. (E-G) Predicted *versus* observed counts (E), proportions (F), and Wrightian fitness (G), respectively. The Wrightian fitness of a strain is defined as the ratio of its proportions at time t+1 and time t. Colours represent the continents (brown: Africa; red: Americas; yellow: Asia; green: Australia; blue: Europe). The dotted line denotes the identity line.

We found that the underlying fitness of a genotype was associated with whether WCV or ACV was being used in that country at that time. *ptxP3*/PRN-positive types, irrespective of *fim3* allele, were the most fit genotypes during the WCV era (relative fitness of 1.11 [95%CI: 1.05-1.17] per year compared to other genotypes circulating at that time, equivalent to a relative fitness of 1.01 [95%CI: 1.00-1.01] per transmission generation). The *ptxP3/fim3-1*/PRN-deficient genotype was the most fit genotype following ACV implementation (relative fitness of 1.30 per year [95%CI: 1.16-1.53], equivalent to a relative fitness of 1.02 per transmission generation [95%CI: 1.01-1.03]) (Figure 4A). We found that the switch from WCV to ACV is associated with an increased fitness of PRN-deficient genotypes, but only in isolates with a *ptxP3* background, with the effect further increased for *fim3-1* genotypes (Figure 4B). On average, PRN-deficient isolates were 1.25 (95%CI: 1.20-1.31) times as fit as PRN-positive isolates following the introduction of ACV (Figure 4C). Where only WCV was used, PRN-deficient isolates were on average 0.94 times (95%CI 0.92-0.96) as fit as their PRN-positive counterparts (Figure 4C). Alternative models that did not allow for changes in fitness following vaccine switch or use a common date across countries for changes in fitness had lower adequacy as measured by Watanabe–Akaike information criterion (Table S3)^21^. Models that allowed for a delay between ACV implementation and fitness change also did not improve model fit (Figure S10, Table S4), suggesting a rapid impact of the vaccine switch on genotype fitness at the population level. Our model was able to recover true fitness parameters when using simulated data with known parameters, even under conditions of biased sampling (Figure S11). Further, our population-level findings are consistent with those from experimental studies. For example, PRN-deficient isolates had increased ability to colonise the respiratory tract of ACV-vaccinated mice than PRN-positive isolates, whereas in unvaccinated control mice, PRN-positive isolates outcompeted PRN-deficient isolates^22^. Mouse models have also shown increased respiratory tract colonization by *ptxP3* than *ptxP1* isolates^23,24^.

**Figure 4:**
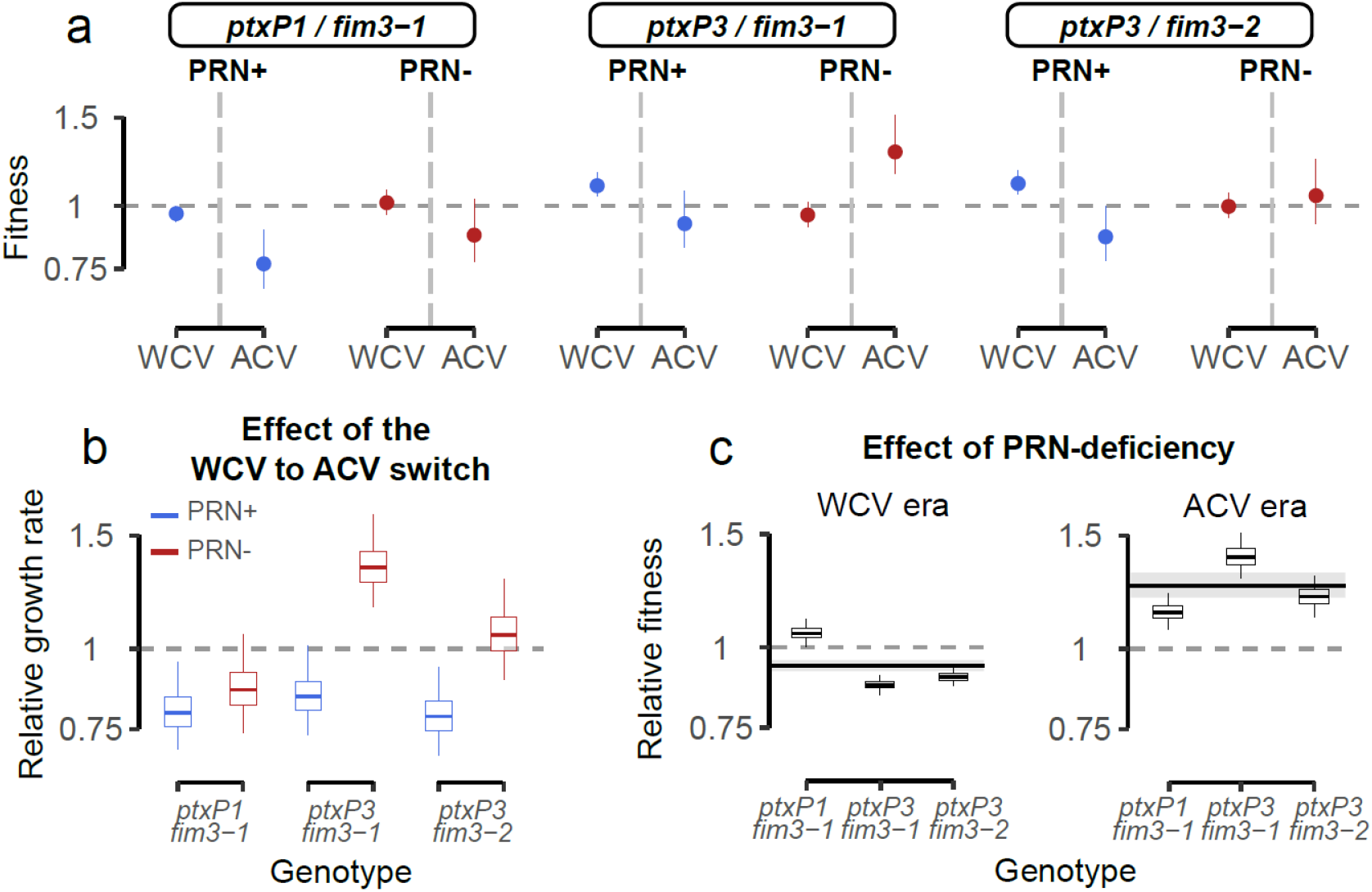
Estimates of fitness of each genotype. (A) Estimated fitness of each genotype as a function of the vaccine era. The dots and lines represent 2.5, 50 and 97.5 percentiles of the posterior distributions. PRN-deficient strains (PRN-) are shown in red, PRN-positive strains (PRN+) are shown in blue. (B) Effect of vaccine switch (whole-cell vaccine [WCV] to acellular vaccine [ACV]) for the different genotypes and PRN expressions. (C) Effect of PRN-deficiency on the different underlying genotypes. Horizontal lines and shaded areas represent the overall mean across all genotypes. The boxplots represent the 2.5, 25, 50, 75, and 97.5 percentiles of the posterior distributions.

The genotype dynamics in Japan represents a key outlier, as the proportion of isolates that were PRN-deficient initially increased following ACV implementation, but unlike most other countries it subsequently decreased after 2005 (Figure 1H)^25^. To date, the main hypothesis to explain this decrease has been a change in ACV formulations to ones that do not contain PRN, which became widely available in 2012^25^. We provide a new explanation for the decrease in PRN-deficient strains. *ptxP1* remained in circulation in Japan after it had largely disappeared elsewhere, and unlike PRN-deficient *ptxP3* strains, PRN-deficient *ptxP1* strains only developed a small fitness advantage post ACV implementation (Figure 4B). The reduction in PRN-deficient strains may simply have been driven by the rise of PRN-positive *ptxP3* strains, which are marginally more fit than *ptxP1* PRN-deficient strains in the ACV context. We expect that this phenomenon can only delay the evolution towards PRN deficiency, as switches to fitter PRN-deficient forms of *ptxP3 B. pertussis* appear inevitable. It remains unclear whether the growing use of ACV formulations without PRN in Japan can further delay this predictable trend.

Our fitness model was unable to capture the genotype dynamics in China, where *ptxP1*-positive strains remain dominant (Figure S9C). This could be explained by the uniquely high prevalence of erythromycin resistant strains^26,27^, which might have a large impact on fitness^28^ but which our model does not take into account.

## Discussion

We have used a large geo-referenced dataset to uncover the spatial spread of an important bacterial threat to human health. We have also identified how vaccine policy decisions can have immediate and far-reaching implications for the fitness of circulating strains, and how small fitness differences at each transmission generation can lead to major changes in lineage composition. Our findings also support and quantify the driving role of acellular vaccines in changing genotype distribution and are consistent with changes in the mutation rate in vaccine antigen genes following ACV implementation^29^. A similar role of vaccine-induced changes in lineage composition has been found for the pneumococcus^30–32^.

As with virtually all phylogenetic studies, the availability of sequences differs substantially by country and year. In particular, our inferences are largely based on sequences from high income countries. WCV remains broadly used in low- and middle-income countries. As we have also quantified genotype fitness when WCV was the dominant vaccine, our findings remain relevant to these undersampled locations, and suggest that, given their increased fitness in WCV environments, *ptxP3*/PRN-positive genotypes will dominate. This is indeed the case for the two countries in our dataset, which have only ever used WCV (Tunisia and Iran). In the event of future switches from WCV to ACV, we could expect PRN-deficient strains to prevail, either through the successful introduction of strains circulating elsewhere or through the loss of PRN expressions in local strains. While this study assesses the impact of vaccine type (WCV or ACV) on strain fitness, increased, sustained sequencing efforts would allow us to tackle more precise questions about whether different WCV or ACV vaccines^33–35^ or vaccination policies and schedules^36,37^ exert distinct pressures. Finally, there may also be other mutations that are linked to changes in fitness that we have not considered here^17^.

This work provides a first detailed view of the strength of spatial structure and rate of geographic spread for a pathogen that is responsible for tens of thousands of deaths each year, highlighting how *B. pertussis* spread is a globally interconnected issue. It further provides a quantitative description of strains fitness and highlights the key role of vaccine policy decision-making in driving ecological change. Multiple co-circulating lineages are a common feature across bacterial disease systems. Our approaches provide an avenue to obtain a much needed mechanistic understanding of how perturbations in population immunity can drive local genetic diversity.

## Methods

### Selection of isolates

We compiled a dataset of 3344 *B. pertussis* whole genome sequences (Table S1 and S2), sampled from a 85-year period (1935-2019) which includes 1011 sequenced French isolates from the National Reference Center (NRC) for Whooping Cough and Other Bordetella Infections in France (Institut Pasteur, Paris, France); 320 newly sequenced European isolates, randomly selected among the 2014-2016 isolates from the *B. pertussis* collections of 12 European countries; and 2013 isolates with high-quality publicly available genomes from 14 countries. The majority of publicly available isolates were generated by the CDC (Atlanta, CDC), who shared medata. Critically, several isolates deposited in NCBI were duplicates from the same isolates; these were removed from the analysis.

### Metadata

We collected metadata for each isolate (date of sampling, continent, country) from linked publications, or NCBI when no publication was listed. The vaccine status of patients was not known. For 3,256 sequences coming from 19 countries, the home district or region of the genome was also recorded. Geographic coordinates were extracted for each location. When the geographic precision was region or district, the coordinates of the most populated city were extracted. We also considered centroids of the whole region as a sensitivity analysis. For each country, we extracted information on vaccine coverage as a function of time (1980-2019, when available) from the Global Health Observatory data repository^38^. We also collected years of implementation of each type of vaccine in the literature. For the acellular vaccine (ACV), we distinguished the year of the first implementation of any ACV (as a booster or a primary vaccination), and the date of ACV introduction for primary vaccination series (Table S1, Figure S8). To maximize statistical power, we focused the analysis on the type of vaccine (WCV, ACV), rather than the specific details of each vaccine (e.g., vaccine provider, vaccine strain used, number of antigens present, presence of a pertactin component). We also chose to focus on the use (or not) of each vaccine in each country, rather than on the country-specific vaccine coverage, assuming that the vaccination coverage is sufficient to exert selective pressure on strains.

### Ethical considerations

The study was coordinated by the French national reference center for pertussis, whose activities are approved by the French supervisory ethics authority (CNIL, n°1474593). Isolates sequenced in this study come from the *B. pertussis* governmental surveillance laboratories of Belgium, Czech Republic, Denmark, Spain, Finland, France, Ireland, Italy, Netherlands, Norway, Sweden and the UK. These isolates were all collected as part of existing public health surveillance approved protocols in each country. No personally identifiable information was used as part of this study.

### Sequencing of the newly sequenced isolates

The study Accession Numbers on the European Read Archive are PRJEB21744, PRJEB42353 and PRJEB45681. Details and accession numbers of the raw sequence data are listed in Table S2. Details on the sequencing protocol can be found in supplementary materials.

### Genomic analysis of PRN-deficient isolates

*De novo* assembly was performed, as previously described^39^. Briefly, paired-end reads were clipped and trimmed (AlienTrimmer^40^), corrected (Musket^41^), merged if needed (FLASH^42^), and subjected to a digital normalization procedure with khmer^43^. For each sample, remaining processed reads were assembled and scaffolded (SPAdes^44^).

We defined the PRN allele of each isolate with BLASTn^45^ using as query a fasta file containing all known PRN alleles.

Next, we defined the pertactin expression (PRN status) of all isolates. As the PRN-deficiency has been shown to be caused by mutations in the promoter or coding regions of the PRN gene^46^, we compiled all the events that cause the PRN deficiency based on the literature and on the analysis of french isolates and gathered all the genomic events identified in PRN-negative isolate in a single fasta file. Then, we assessed the PRN status of all isolates with BLASTn^45^ using as query the fasta file (Table S5, File S1). In addition, we used IS_mapper/0.1.5.1 from fastq files looking for IS*481* from Tohama (BP0080), which is the main insertion element in *B. pertussis*^*47*^ that can insert within the prn gene or promoter. We made use of 839 french isolates for which we have Western blots available^2^, and checked the correspondence between genomics and experimental results. In our dataset, we identified 1722 (51.5%) PRN-positive isolates, 1472 (44.0%) PRN-deficient and 150 (4.5%) isolates with unknown status.

### Nucleotide Polymorphism variations (SNP) detection

SNP detection was conducted using an in-house pipeline available on GitHub. Briefly, adapters and barcodes were stripped from the fastq data and the reads were quality filtered and trimmed using a Phred quality threshold score of 30 using Cutadapt^48^. We checked the quality of each fastq file using FastQC^49^. Reads were mapped against the complete Tohama I reference genome (Accession number: NC_002929, using BWA-MEM algorithm^50^. Extraction of SNP was achieved with the GATK HaplotypeCaller, with ERC GVCF settings^51^. We then built an in-house filtering script using R^52^. We kept variants for which the Phred quality score was higher than 30, with a minimum read depth of 5, with at least 2 reads in both the forward and reverse directions. We called a position a ‘N’ when <40% of the reads were different from the reference, we used the IUPAC code for positions with between 40% and 80% of reads not matching the reference, and we called variants only for positions with >80% reads not matching the reference. Moreover, to improve homogeneity across the dataset across the world, we removed positions in the alignment where 25% of the bases were ‘N’. Further, we filtered out repeated regions (IS481, IS1002 and IS1663^47^), and phage regions using Phaster^53^. We also checked for recombination in our alignment using Gubbins^54^. As a result, we obtained an alignment of 8,105 SNPs, consistent with other publications^17^.

### Genotyping *ptxA, ptxP, fim2 and fim3*

We genotyped all isolates for the genes pertussis toxin A (*ptxA*), pertussis toxin promoter (*ptxP*), type 2 fimbrial protein (*fim2*) and type 3 fimbrial protein (*fim3*) using BLAST^45^. The sequences used as references for the alleles *ptxA, ptxP, fim2* and *fim3* are available in GitHub (https://github.com/noemielefrancq/GlobalPhylogeographyPertussis).

### Phylogenetic analysis

*B. pertussis* strain genotyping has been used to reconstruct the evolutionary and spatial history of observed infections, however, previous efforts have relied on pre-genomics markers or individual country data^27,29,39,55–59^. In this project we used whole genome sequences and combined data across multiple countries.

We used the SNP-based alignment to reconstruct the phylogenetic relationships of the isolates. We built maximum-likelihood trees using IQ-tree^60^. TempEst was used to check for temporal signal in the data (Figure S12)^61^. Maximum Clade Credibility (MCC) trees were inferred using a Markov Chain Monte Carlo (MCMC) Bayesian approach implemented on the program BEAST 1.10.4^62^, under a GTR substitution model^63^ accounting for the number of constant sites, a relaxed lognormal clock model^64^ and a skygrid population size model^65^. Three independent Markov chains were run for 300,000,000 generations each, with parameter values sampled every 10,000 generations. Runs were optimized using the GPU BEAGLE library^66^. Chains were manually checked for convergence (ESS values > 200) using the Tracer software^67^. We manually removed a 20% burnin.

Furthermore, we used a discrete model attributing state characters representing the isolation country of each strain with the Bayesian Stochastic Search Variable (BSSVS) algorithm^68^, implemented in BEAST. This method estimates the most probable state at each node in the MCC trees, allowing us to reconstruct plausible ancestral states (here, country) on these nodes. Because of the large nature of our dataset (3344 genomes from 23 countries, with 8925 variable positions) running this model would have been computationally challenging. Thus, we ran the model on a subset of trees (N=10,000) extracted from the posterior distribution of trees generated by the initial BEAST run. We ran three independent Markov chains for 5,000,000 generations each, with parameter values sampled every 1,000 generations. We manually checked for convergence using the Tracer software^67^ and removed a 10% burnin. We used treeannotator to summarize the posterior trees into a Maximum Clade Credibility (MCC) tree. The full MCC tree is presented in Figure S13.

### Expansion rate of *B. pertussis*

We assume that the sequences in our dataset are representative of what is circulating in that location and at that time. In spatially structured transmissions, as pathogens spread away from each other, we would expect there to be increased spatial distance between cases as they become separated by more and more transmission events^20^. To explore whether this occurs, we compared evolutionary time between *B. pertussis* bacteria with their spatial separation. For each location considered, pairs of bacteria are grouped by the evolutionary time that separates them based on the time resolved phylogeny. For each group of bacteria pairs, we then compute the mean spatial distance that separates them. We reconstruct 95% confidence intervals using bootstrap. We resample all the bacteria with replacement, allowing for even sampling by location, over 200 resampling events and recalculate the mean distance that separates each group of bacteria pairs each time. The 95% confidence intervals are the 2.5% and 97.5% quantiles from the resultant distribution. We separately consider pairs coming from the same country (France, Japan, US). This list of countries was selected as they represent the locations with most sequences available. We separately repeated this analysis using pairs of sequences from across Europe.

### Estimating the probability that a pair of cases are from the same transmission chain

We consider a pair of cases that came within the same year to be part of the same transmission chain if their most recent common ancestor (MRCA) was within the past two years of the earlier of the two cases. We choose this cutoff as two years represents a recent introduction into apopulation, within which pairs are likely to remain transmission related. We explore the sensitivity of this cutoff (Figure S14).

We can derive an expression for the probability *P*_*x*_ that a pair of *B. pertussis* cases are from the same transmission chain within a location *loc*^20^:

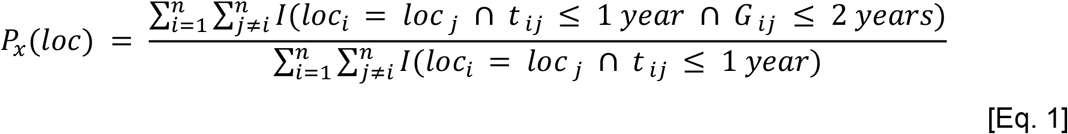

where *n* represents the number of bacteria for which sequence data are available, *t*_*ij*_ is the time between the cases, and *G*_*ij*_ is the time to the MRCA from the earlier of *i* and *j. I*is an indicator function.

### Estimation of effective number of transmission chains

The reciprocal of *P*_*x*_*(loc)* is an estimate of the size of the pool of discrete transmission chains *η(loc)* that infect pairs of individuals separated within a location *loc*. It also represents the lower limit of the number of chains circulating within a location *loc*, as previously shown^20^.

We reconstruct 95% confidence intervals using bootstrap. We resample all the bacteria with replacement, allowing for even sampling by location, over 200 resampling events and recalculate the statistic each time. The 95% confidence intervals are the 2.5% and 97.5% quantiles from the resultant distribution.

### Effective number of transmission chains for different population sizes in regions

We explore whether the effective number of chains circulating within regions depends on the size of the population of this region, using Equation 1. We estimate the mean probability *P*_*x*_*(loc, pop*_*1*_, *pop*_*2*_) that a pair of *B. pertussis* cases within the same region, with a population size between ***pop***_***1***_ and ***pop***_***2***_ are from the same transmission chain:

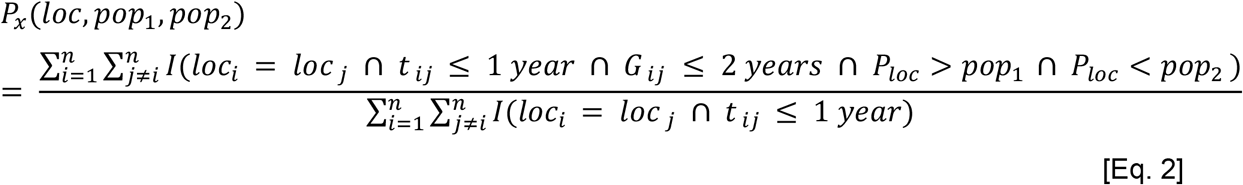

The effective number of transmission chains *η(loc, pop*_*1*_, *pop*_*2*_) is then given by the reciprocal of *P*_*x*_*(loc, pop*_*1*_, *pop*_*2*_).

We reconstruct 95% confidence intervals using bootstrap, as detailed above.

In a sensitivity analysis, we considered different MRCA cutoffs (1 and 3 years) to define transmission chains, and obtained similar results (Figure S14).

### Relative risk that a pair of bacteria have a MRCA within a defined period, when coming from the same location, versus different locations

To better understand the spread of *B. pertussis* within districts in France, countries in Europe, states in the US and between continents, we characterize the similarity in bacteria within these spatial scales relative to that observed between spatial scales. In each case, we estimate *RR*_*loc*_*(g*_*1*_, *g*_*2*_), the probability that a pair of bacteria *within a location loc* that were isolated within the same year of each other had an MRCA within range having an MRCA within *g*_*1*_-*g*_*2*_ relative to the probability that a pair of bacteria from different locations, that were isolated within the same year of each other, had an MRCA within that particular range^20^. For the range *g*_*1*_-*g*_*2*_, we used sliding windows of time going from 0 years to 20 years.

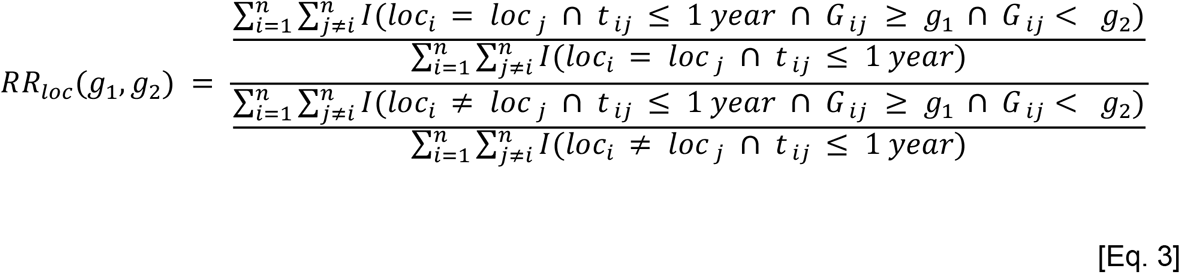

By conditioning on spatial and temporal location of sequences, this approach minimizes the impact of underlying sampling biases in which cases were sequenced. We reconstruct 95% confidence intervals using bootstrap, as detailed above.

### Estimating the fitness of *B. pertussis* strains

To quantify the fitness of *B. pertussis* strains, we develop a model that makes use of the isolates from 20 countries with more than 10 sequences in our dataset (fewer sequences than this threshold did not allow for sufficient data points to provide a robust estimate of genotype distributions over time). First, we assign all tips and nodes in the tree a strain type from one of six possible types based on PRN, *ptxP* and *fim3* status. We consider only isolates for which the *ptxP3* allele was 1 or 3, the *fim3* allele was 1 or 2, and the PRN expression was known. While there exists other genotypes, these are rare and most of the other types, for example the ptxA alleles, almost completely overlap with the *ptxP* alleles. Assigning the genotypes *ptxP1* versus *ptxP3*, and *fim3-1* versus *fim3-2* is straightforward as they are all monophyletic^17^. However, the PRN-deficiency is a highly homoplasic trait, so assigning a PRN expression to each node is more complicated. Briefly, we know ancestral *B. pertussis* sequences all produce PRN, and that the deficiency appears by mutations in the promoter or coding regions of the PRN gene^46^. Moreover, because of the complexity of the mutations, reversions are highly unlikely, and have to our knowledge never been reported. Thus, the PRN expression of each node can be easily reconstructed by finding all the monophyletic clades for which isolates are PRN-deficient. To do this, we use an in-house algorithm (script available in the GitHub).

We separately assign a country to each node and tip. Country information for the nodes was extracted from the BEAST discrete reconstruction presented previously. In order to maximize the correct country assignment for nodes, we only consider nodes for which the country’s posterior probability was >0.9. In addition, we excluded nodes that were distant by more than 5 years from any tip.

Next, we compute *f*_*i,ref*_, the relative abundance of each strain *i* with respect to a chosen *ref* strain. We chose the *ptxP3*/*fim3-1*/PRN-positive genotype as the reference strain. In a sensitivity analysis, we found that the choice of the reference strain did not affect the results, providing that the chosen genotype was present in all countries, for most of the years. We then use a simple logistic model to capture the evolution of this abundance, at each time t:

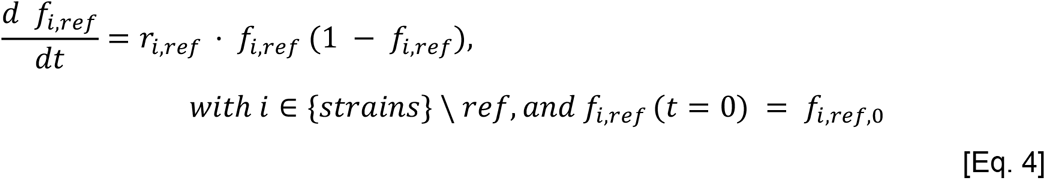

where *r*_*i,ref*_ is the growth rate of that abundance, shared across all countries and *f*_*i,ref,0*_ is the initial relative abundance of the strain *i* with respect to a chosen *ref*. To control for the varying presence of all circulating strains through time, we present fitness as the *average relative growth rate* <u>*r*_*i*_</u> for each strain *i* with respect to randomly selected strain in the population:

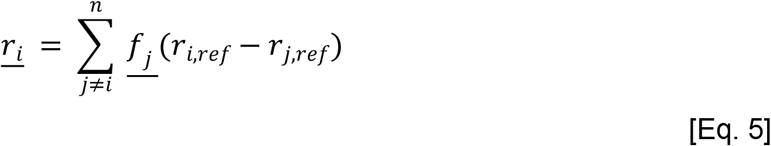

where *n* in the number of strains, and *f*_*j*_ is the average frequency of strain *j*on the period of time considered.

This *average relative growth rate* <u>*r*_*i*_</u> can be identified as the selection rate coefficient *s* of the strain *i* in the population considered^69–71^.

We can further multiply the selection rate by the mean generation time *T* to obtain the dimensionless selection coefficient *s*_*T*_, the relative fitness advantage per transmission generation^70^:

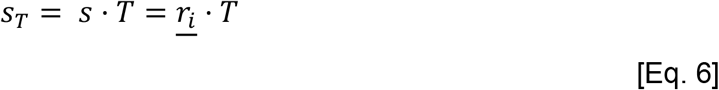

We use a mean generation interval of 22.8 days (95%CI: 22.1-23.5) to compute *s*_*T*_^72^.

The selection coefficients *s* and *s*_*T*_ represent one of the most possible direct measures of the fitness advantage of any new variant, and are the best possible predictors of whether or not it is expected to increase in frequency during an outbreak^71,73^.

We fit this model to all countries-specific time series in our dataset with the Rstan package^74^, using a Negative Binomial likelihood with an overdispersion parameter *δ*. We fit the frequencies as simplex vectors for each country, with the oldest strain frequency, *f*_*1,0*_, having a *normal(1, 0*.*1)* prior, based on a previous global study^17^. We use *cauchy(0, 0*.*15)* for the relative growth rate parameters *r*_*i,ref*_. We ran this model on 3 independent chains with 2,000 iterations and 50% burn-in. We use 2.5 and 97.5 quantiles from the resulting posterior distributions for 95% credible intervals of the parameters.

### Investigating changes in *B. pertussis* fitness across vaccine eras

To investigate whether *B. pertussis* strain fitness changed across vaccine eras, we compiled dates of vaccination changes (Table S1) and used our fitness model and estimate *r*_*i*_ on:

- WCV era, defined as the period from the WCV implementation in each country, to the first ACV implementation (as a booster or primary vaccine);
- the ACV era, defined as the period from the first ACV implementation to now.

In a sensitivity analysis, we consider a range of different definitions for these eras, including the mean implementation year of the WCV and ACV across countries, and start at different vaccine coverages (Table S3). We also considered models without a vaccine switch. Model comparison was done using the Watanabe–Akaike information criterion (WAIC) implemented in the *loo* package^21,75^. Estimated from this sensitivity analysis are presented in Figure S15.

Additionally, to test the predictive power of our model, a held out analysis was performed. We held out 10% of the country-year data from the model fitting process, and compared the prediction with the actual observed values (Figure S16).

We developed a simulation framework where the true growth rates parameters were known, and assessed the performance of our model to estimate the fitness of different strains in a population (see Supplementary materials).

## Acknowledgments

We thank Dr. C. Schot for her contribution to isolates and genomic data collection. We thank Gabriele Buttinelli and Luigina Ambrosio (Dept. of Infectious Diseases, Istituto Superiore di Sanità, Rome, Italy) and Carlo Concato (Virology Unit, Bambino Gesù Children’s Hospital, IRCCS, Rome, Italy) for their contribution to Italian isolates collection and preparation. We thank Dr. M. Weigand and his team at CDC (Atlanta, USA) for making sequences publicly available and sharing metadata. The study was supported financially by the French Government Investissement d’Avenir grant ANR-16-CONV-0005 (INCEPTION project). The National Reference Center for Whooping Cough and Other Bordetella Infections receives support from Institut Pasteur and Public Health France (Santé publique France, Saint Maurice, France). This work was also supported financially by the French Government’s Investissement d’Avenir program Laboratoire d’Excellence “Integrative Biology of Emerging Infectious Diseases” (ANR-10-LABX-62-IBEID) and a European Research Council (No. 804744). PL acknowledges funding from the European Research Council under the European Union’s Horizon 2020 research and innovation programme (grant agreement no. 725422-ReservoirDOCS).

## Author’s contributions

J.T., S.C., H.S. and S.B. conceived the study. N.L. and H.S. developed the methods and performed the modelling analyses, supported by S.L.H., P.L., S.B. and J.P. V.B., N.L and N.F. performed the genomic analyses. N.L., V.B, N.F., A-M.B., T.B., T.A., K.F., D.F.V., N.K.F., J.G.L., K.G., A.H., Q.H., D.L., H.M., D.P., P.S., J.Z, S.G., J.T., and S.B. contributed to isolates and genomic data collection. V.B., N.A., A.L., S.G., T.B., P.S., H.M. and M.S. sequenced the isolates. N.L. and V.B. had access to and verified all the data. N.L., H.S. and S.B. wrote the first draft of the manuscript. All authors provided input to the manuscript and reviewed the final version.

## Additional Information

Supplementary Information is available for this paper. Code will be made available at: https://github.com/noemielefrancq/GlobalPhylogeographyPertussis/. All sequences generated within this study were deposited in ENA (Table S2 accession numbers)[will be made available on acceptance of the manuscript]. The metadata attached to each sequence is available in Table S2. Correspondence should be addressed to Dr. Sylvain Brisse and Dr. Henrik Salje.

