## Supplementary information for "Spatial dynamics and vaccine-induced fitness changes of *Bordetella pertussis*"

### Supplementary methods

##### DNA extraction and Illumina NextSeq Sequencing for the newly sequenced isolates

Isolates were grown at 36 °C for 72 h on Bordet-Gengou Agar (Becton Dickinson, Le Pont de Claix, France) supplemented with 15% defibrinated horse blood (Biomerieux, Marcy l'Etoile, France) and sub-cultured in the same medium for 24 h. Bacteria were suspended in physiological water to reach $OD_{650}$ of 1 and 400 ml were pelleted. Pellets were suspended in 100 ml of PBS 1X, 100 ml of lysis buffer (Roche) and 40 ml of proteinase K, heated at 65 °C for 10 minutes, then at 95 °C for 10 minutes and further used for DNA extraction. Libraries were constructed using the Nextera XT DNA Library Preparation kit (Illumina, Inc., San Diego, CA) and whole genome sequencing was performed on a NextSeq 500 system (Illumina, USA) using a 2×150 paired-end protocol at the Mutualized Platform for Microbiology of Institut Pasteur. A minimum average read depth of 50X was expected.

##### Simulation study to demonstrate the robustness of the fitness model

To assess the performance of our model to estimate the fitness of different strains in a population, we developed a simulation framework where the true growth rates parameters were known.

For a period of 100 years, we simulate an epidemic, seeded by 10 infections from two different strains each, where the number of cases grows exponentially each day. We specified the characteristics of the two strains (Figure S10A):

- Strain 1: initial proportion $f_{0,1} = 0.95$, and $R_{0,1} = 1.1$;
- Strain 2: initial proportion $f_{0,2} = 0.05$, and $R_{0,2} = 1.2$.

Thus, setting a common generation interval T for both strains, the selective advantage (fitness) of the strain 1 in the population is:

$$s_{T,1} = log(\frac{R_{0,1}}{R_{0,2}})$$

[Eq. 10]

We then sample from this epidemic N=2000 infected individuals, with two different strategies (Figure S10B):

- uniform sampling throughout this epidemic: 20 sequences per year;
- biased sampling: the first 10 years were not sampled, years 11-40 were very sparsely sampled (1-2 sequences per year), then the following years were sampled with a

minimum of 10 sequences, and as high as 90 sequences some years. This procedure is meant to closely mimic the temporal sampling structure of our study.

We use the simulated data to estimate the relative fitness of strain 1 compared to strain 2 using our fitness model and compared with the known true values (Figure S10C-E) .

#### Supplementary tables

Table S1: Summary of the countries represented in the dataset, and their vaccine implementation dates.

Table S2: Description of the 3344 isolates used in this study.

Table S3: Comparison of different fitness models.

Table S4: Comparison of the fitness models allowing a delay between ACV implementation and fitness change.

Table S5: Description of the different events impairing *prn* gene expression.

#### Supplementary figures

Figure S1: Median spatial distance between worldwide *B. pertussis* pairs separated by different short evolutionary times.

Figure S2: Sensitivity analysis for the definition of population centroids.

Figure S3: Proportion of cases within different MRCA windows, across locations.

Figure S4: Proportion of pairs from the same transmission chains and number of effective transmission chains within regions, as a function of population size.

Figure S5: Number of effective transmission chains for different areas and population densities.

Figure S6: Risk ratio that a pair of bacteria have a MRCA within a defined period, when coming from the same versus different district, for different countries.

Figure S7: Risk ratio that a pair of bacteria have a MRCA within a defined period, when coming from the same versus different continent, for different pairs of continents.

Figure S8: Implementation of whole-cell and acellular vaccines by country.

Figure S9: Model fits for all countries.

Figure S10: Model fit for different delays between ACV implementation and fitness change.

Figure S11: Results of the simulation study for our fitness model, with or without biased sampling.

Figure S12: Temporal signal in the dataset.

Figure S13: Maximum clade credibility tree for all the isolates.

Figure S14: Sensitivity of transmission chains estimates to changing MRCA cutoff.

Figure S15: Sensitivity analysis: estimates with a range of different models.
Figure S16: Held out fitness model.

#### Supplementary data

https://github.com/noemielefrancq/GlobalPhylogeographyPertussis [will be online upon publication]

**Table S1: Summary of the countries represented in the dataset, and their vaccine implementation dates.**

We report Whole-Cell Vaccine (WCV), first and primary series Acellular Vaccine (ACV) implementation years for countries with at least 10 isolates (NA: not applicable).

| Country | Number sequences | Collection years | Study of genomic sequences | First WCV implementation[a] | First ACV implementation[b] | ACV implementation as primary vaccination | Reference for vaccination dates |
|---|---|---|---|---|---|---|---|
| **Australia** | 102 | 1997-2017 | NCBI, [1] | 1942 | 1997 | 1999 | [2,3] |
| **Belgium** | 30 | 2014-2016 | This study | 1950 | 1999 | 1999 | [4] |
| **Brazil** | 1 | 2011 | NCBI | NA | NA | NA | NA |
| **Canada** | 40 | 2012-2013 | [5] | 1943 | 1997 | 1997 | [6] |
| **China** | 49 | 1951-2015 | [7] | 1960 | 2007 | 2012 | [8,9] |
| **Czech Republic** | 24 | 2014-2016 | This study | 1958 | 2007 | 2007 | [10] |
| **Denmark** | 16 | 2014-2016 | This study | 1961 | 1997 | 1997 | [4] |
| **Spain** | 30 | 2014-2016 | This study | 1965 | 1999 | 2005 | NA |
| **Finland** | 30 | 2014-2016 | This study | 1952 | 2003 | 2005 | [4] |
| **France** | 1011 | 1953-2019 | This study, [11–14] | 1959 | 1998 | 2004 | [4] |
| **Guatemala** | 6 | 2014 | NCBI | NA | NA | NA | NA |
| **Ireland** | 42 | 2012-2016 | This study | 1952 | 1996 | 1996 | [15] |
| **Israel** | 18 | 2006-2011 | [16] | 1957 | 2002 | 2002 | [17] |
| **Iran** | 53 | 2008-2016 | [18] | 1950 | Not implemented | Not implemented | [18] |

| | | | | | | | |
|---|---|---|---|---|---|---|---|
| **Italy** | 29 | 2014-2016 | This study | 1961 | 1995 | 1995 | [4] |
| **Japan** | 190 | 1954-2014 | [19] | 1947 | 1981 | 1981 | [20] |
| **Mexico** | 4 | 2008-2014 | NCBI | NA | NA | NA | NA |
| **Netherlands** | 30 | 2014-2016 | This study | 1953 | 2001 | 2005 | [4] |
| **Norway** | 30 | 2014-2016 | This study | 1952 | 1998 | 1998 | [4] |
| **Sweden** | 30 | 2014-2016 | This study | 1953 | 1996 | 1996 | [4] |
| **Tunisia** | 10 | 2014-2014 | [21] | 1979 | Not implemented | Not implemented | [22] |
| **UK** | 112 | 1967-2016 | This study, [23] | 1957 | 2000 | 2000 | [4] |
| **US** | 1457 | 1935-2019 | [24–28], NCBI | 1948 | 1992 | 1997 | [29,30] |

[a] Some countries removed the WCV before implementing the ACV, such as Sweden between 1979 and 1996.

[b] The first ACV implementation date takes into account any implementation of an ACV (booster or primary vaccination).

**Table S2: Description of the 3344 isolates used in this study.**

Attached Excel file

**Table S3: Comparison of different fitness models.**

*Primary ACV* refers to the implementation of ACV as primary vaccination. *Any ACV* refers to the first ACV implementation date, either as a primary vaccination or booster. Model adequacy has been measured with the Watanabe–Akaike information criterion[31].

| Model | Start of vaccine pressure | Shift in fitness | WAIC | Difference to the best model | pWAIC |
|---|---|---|---|---|---|
| **1** | 1955 | 2000 | 1834.91 | 42.07 | 41.20 |
| **2** | 1955 | Primary ACV | 1821.21 | 28.37 | 40.75 |
| **3** | 1955 | Any ACV | 1795.32 | 2.48 | 39.81 |
| **4** | 1955 | None | 1954.89 | 162.05 | 37.93 |
| **5** | 80% coverage | 2000 | 1840.99 | 48.15 | 42.65 |
| **6** | 80% coverage | Primary ACV | 1832.87 | 40.03 | 41.64 |
| **7** | 80% coverage | Any ACV | 1809.81 | 16.97 | 40.96 |
| **8** | 80% coverage | None | 1933.33 | 140.49 | 38.81 |
| **9** | 90% coverage | 2000 | 1846.62 | 53.78 | 42.63 |
| **10** | 90% coverage | Primary ACV | 1839.30 | 46.46 | 41.80 |
| **11** | 90% coverage | Any ACV | 1814.32 | 21.48 | 41.41 |
| **12** | 90% coverage | None | 1940.48 | 147.64 | 38.75 |
| **13** | WCV | 2000 | 1809.33 | 16.49 | 40.49 |
| **14** | WCV | Primary ACV | 1810.04 | 17.20 | 40.76 |
| **15** | **WCV** | **Any ACV** | **1793.28** | **0.44** | **39.88** |
| **16** | WCV | None | 1942.91 | 150.07 | 36.80 |

**Table S4: Comparison of the fitness models allowing a delay between ACV implementation and fitness change.**

In each model, we allow for a delay of *+/- X years* between the first ACV implementation (denoted *Any ACV +/- X years*) and the fitness change. Model adequacy has been measured with the Watanabe–Akaike information criterion[31].

| Model | Start of vaccine pressure | Shift in fitness | WAIC | Difference to the best model | pWAIC |
|---|---|---|---|---|---|
| **15_m10** | WCV | Any ACV -10y | 1802.29 | 10.51 | 39.81 |
| **15_m9** | WCV | Any ACV -9y | 1800.72 | 8.94 | 40.03 |
| **15_m8** | WCV | Any ACV -8y | 1798.60 | 6.82 | 39.50 |
| **15_m7** | WCV | Any ACV -7y | 1799.15 | 7.37 | 40.32 |
| **15_m6** | WCV | Any ACV -6y | 1797.12 | 5.34 | 39.86 |
| **15_m5** | WCV | Any ACV -5y | 1795.90 | 4.12 | 39.88 |
| **15_m4** | WCV | Any ACV -4y | 1795.26 | 3.48 | 40.05 |
| **15_m3** | **WCV** | **Any ACV -3y** | **1792.32** | **0.54** | **39.18** |
| **15_m2** | **WCV** | **Any ACV -2y** | **1792.59** | **0.81** | **39.57** |
| **15_m1** | **WCV** | **Any ACV -1y** | **1791.78** | **0.00** | **39.44** |
| **15** | **WCV** | **Any ACV 0y** | **1793.28** | **1.50** | **39.88** |
| **15_p1** | **WCV** | **Any ACV 1y** | **1792.84** | **1.06** | **39.56** |
| **15_p2** | **WCV** | **Any ACV 2y** | **1794.07** | **2.29** | **39.87** |
| **15_p3** | WCV | Any ACV 3y | 1795.27 | 3.49 | 39.75 |
| 15_p4 | WCV | Any ACV 4y | 1795.32 | 3.54 | 39.38 |
| **15_p5** | WCV | Any ACV 5y | 1798.38 | 6.60 | 40.18 |
| **15_p6** | WCV | Any ACV 6y | 1798.22 | 6.44 | 39.72 |
| **15_p7** | WCV | Any ACV 7y | 1799.02 | 7.24 | 39.86 |
| **15_p8** | WCV | Any ACV 8y | 1800.30 | 8.52 | 39.74 |
| **15_p9** | WCV | Any ACV 9y | 1802.40 | 10.62 | 39.96 |
| **15_p10** | WCV | Any ACV 10y | 1804.35 | 12.57 | 39.63 |

**Table S5: Description of the different events impairing *prn* gene expression.**

Attached Excel file

**Figure S1: Median spatial distance between worldwide *B. pertussis* pairs separated by different short evolutionary times.**

Here we present a global average of the relationship between evolutionary distance and spatial distance for short evolutionary times (0 to 6 years) using all isolates with geolocated information. The shaded area represents 95% CIs.

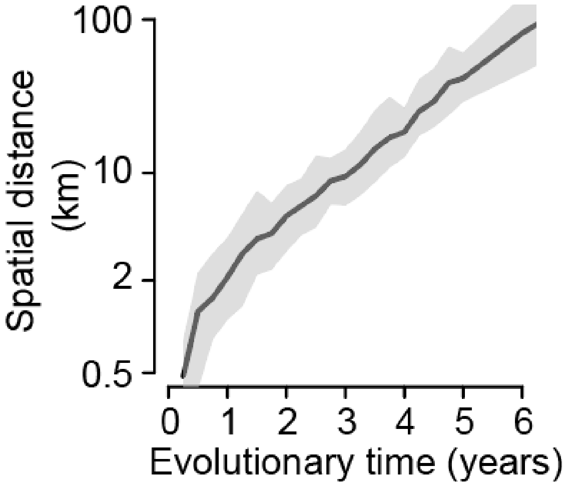

**Figure S2: Sensitivity analysis for the definition of population centroids.**

(A) Comparison of the distances between *B. pertussis* pairs of sequences in the US, computed either by taking the coordinates of the most populated city in each state (x-axis) or the population center of each state (y-axis). (B-C) Median spatial distance between *B. pertussis* pairs, using either (B) the most populated city coordinates, or (C) the population center of each state, separated by different evolutionary times.

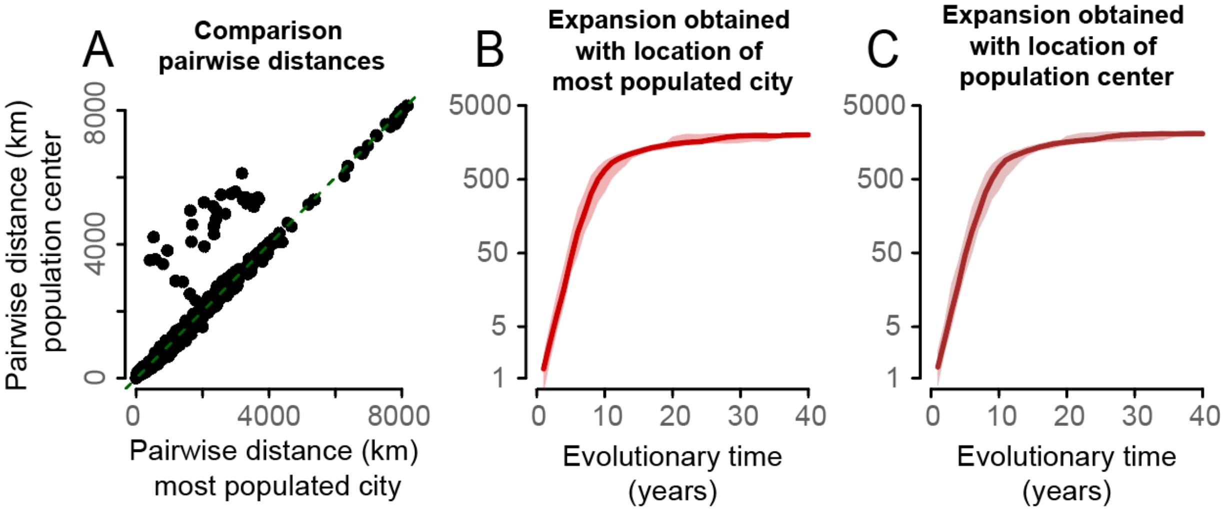

**Figure S3: Proportion of cases within different MRCA windows, across locations.**

**(A)** Breakdown of the MRCA separating pairs of *B. pertussis* isolated within the same year, within districts, for 12 European countries, Iran and Japan. **(B)** Same as (A), but considering isolates within the same countries, irrespective of the district, for 9 European countries (Belgium, Italy and Spain were excluded as only one district was represented for each of them). **(C)** Same as (A), but considering isolates within the same continent, for the Americas, Asia and Europe.

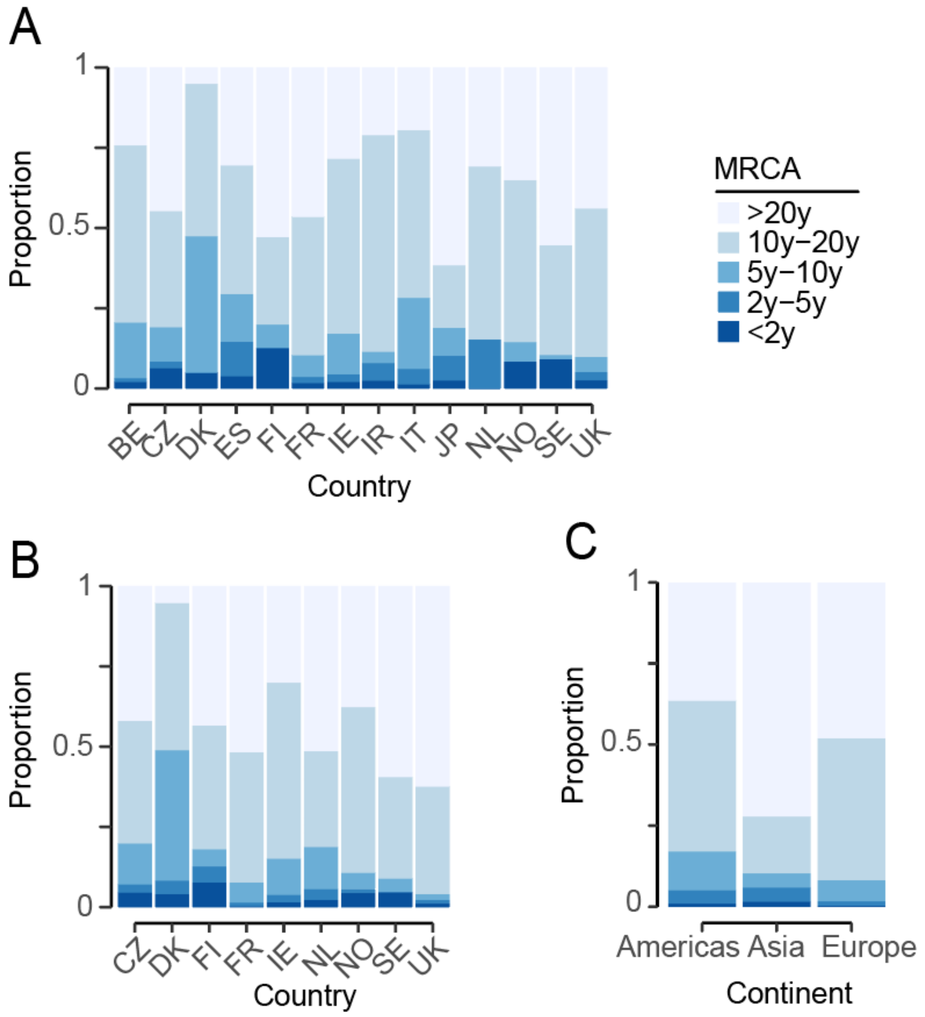

**Figure S4: Proportion of pairs from the same transmission chains and number of effective transmission chains within regions, as a function of population size.**

**(A-C)** Proportion of pairs within a region that belong to the same transmission chain (defined as MRCA<2y), as a function of population size (average from 19 countries), by continent. Proportions are computed for rolling windows of population sizes. Dashed line represents model fit assuming an exponential relationship between the two and the grey shaded region 95% confidence intervals. **(D-F)** Number of transmission chains within regions, as a function of the population size, by continent. The numbers are computed for rolling windows of population sizes. Dashed line represents model fit and the grey shaded region 95% confidence intervals.

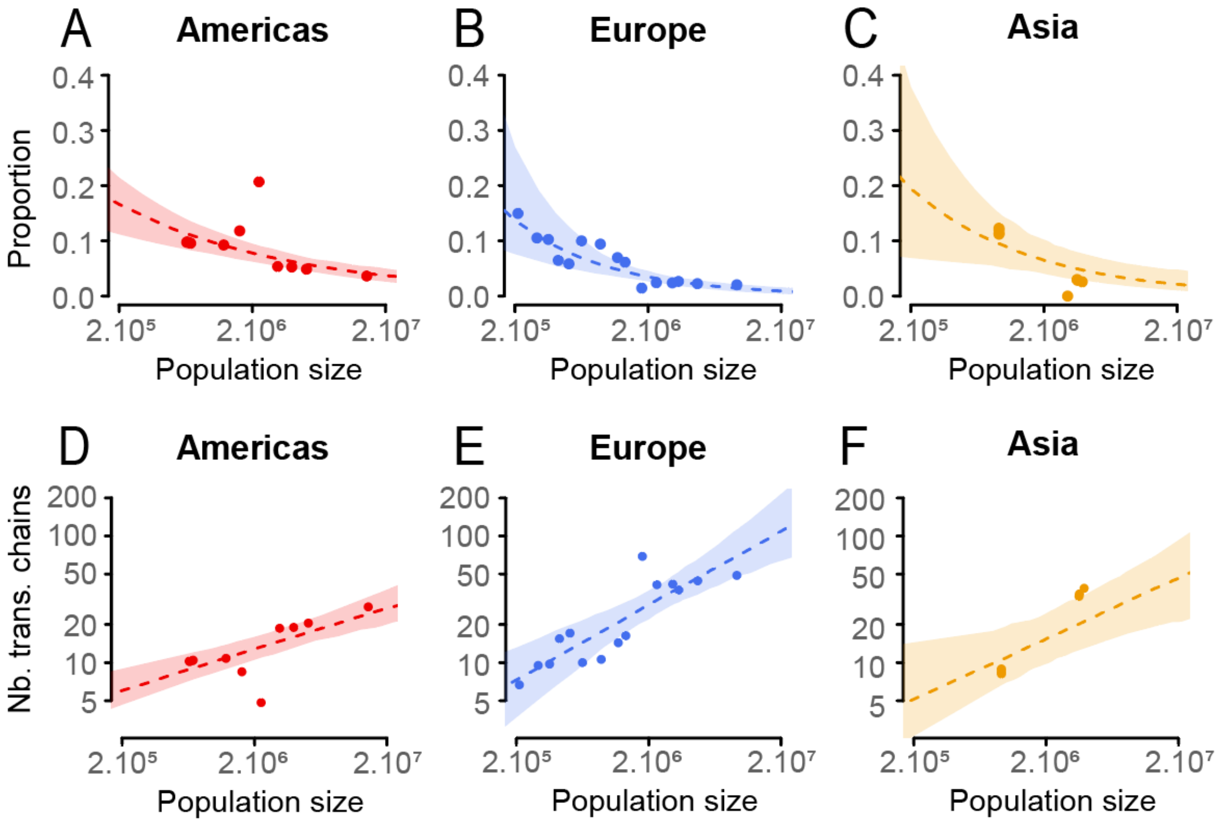

**Figure S5: Number of effective transmission chains for different areas and population densities.**

**(A)** Number of transmission chains within regions, as a function of the area. Dashed line represents model fit and the grey shaded region 95% confidence intervals. **(B)** Same as (A), but as a function of population density. The numbers of transmission chains are computed for rolling windows of area (A) or density (B).

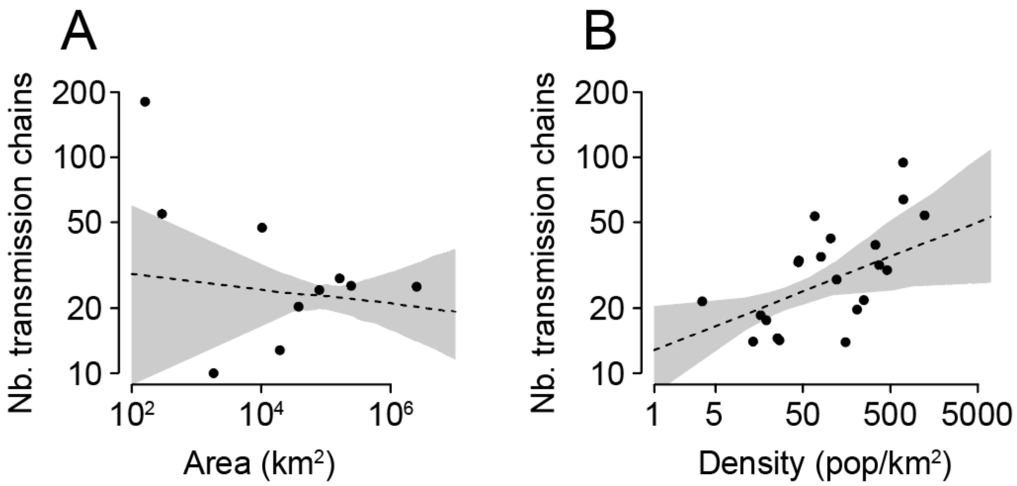

**Figure S6: Risk ratio that a pair of bacteria have a MRCA within a defined period, when coming from the same versus different district, for different countries.**

Risk ratio that a pair of bacteria have a MRCA within a defined period, when coming from the same versus different district in China **(A)**, France **(B)**, Japan **(C)** and UK **(D)**, for which we have enough data available (minimum of 50 sequences per country, with at least 2 districts represented). The shaded regions represent 95% confidence intervals.

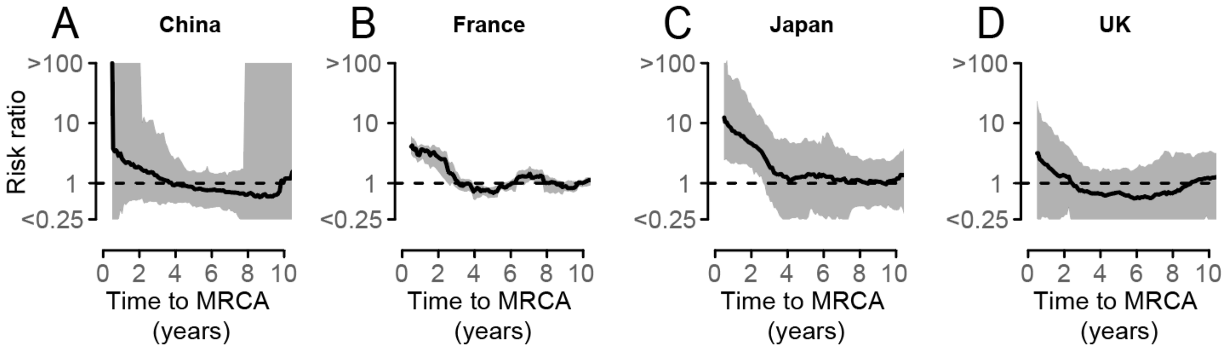

**Figure S7: Risk ratio that a pair of bacteria have a MRCA within a defined period, when coming from the same versus different continent, for different pairs of continents.**
Risk ratio that a pair of bacteria have a MRCA within a defined period, when coming from the same versus different continent, different pairs of continents: Americas and Australia **(A)**, Americas and Asia **(B)**, Americas and Europe **(C),** Asia and Australia **(D)**, Asia and Europe **(E)**, and Australia and Europe **(F)**. The shaded regions represent 95% confidence intervals.

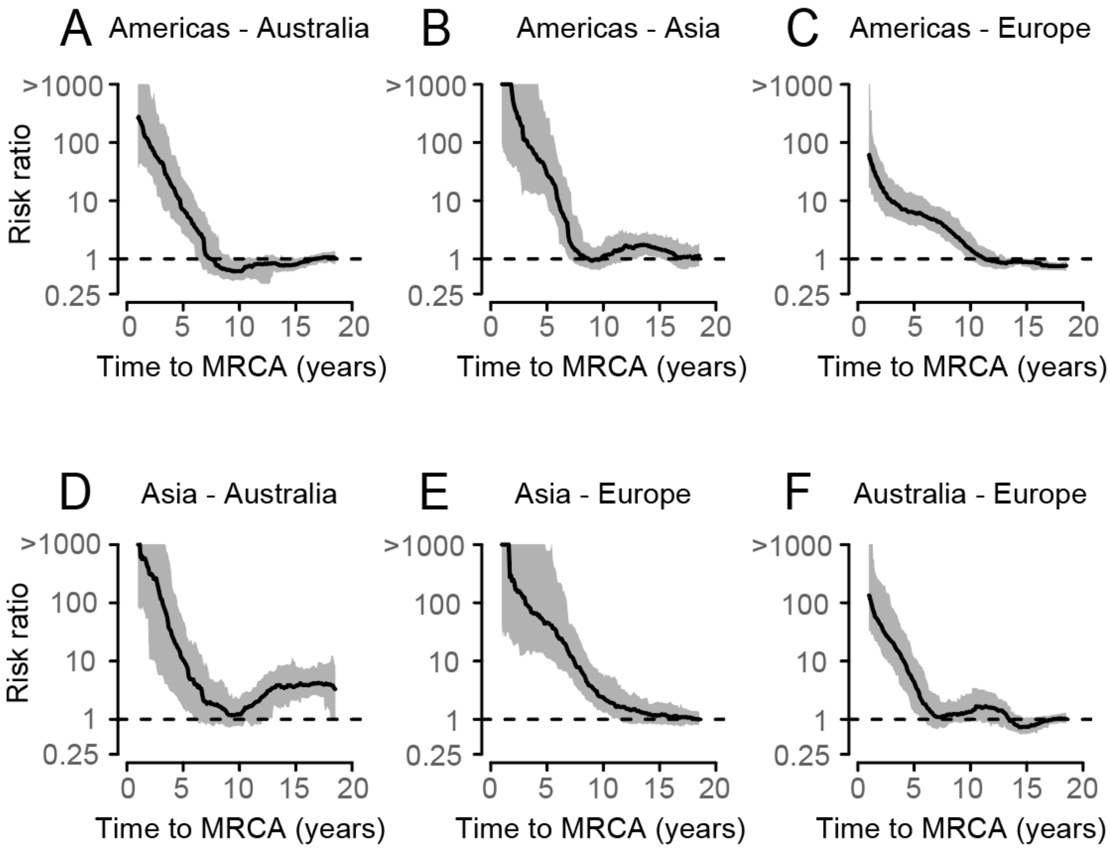

**Figure S8: Implementation of whole-cell and acellular vaccines by country.**

For each country, we plot the type of vaccines that was implemented by a line of different color; grey: no vaccine; blue: whole-cell vaccine (WCV); green: acellular vaccine (including when only as a booster, or in only part of the country); orange: primary acellular vaccine (ACV). Blue squares represent the years of implementation of the WCV for each country, green diamonds show the first implementation of any acellular vaccine, and orange triangles show the implementation of the ACV as primary vaccination. Some countries directly implemented ACV as the primary, country-wide vaccination. Further details and references are shown in Table S1.

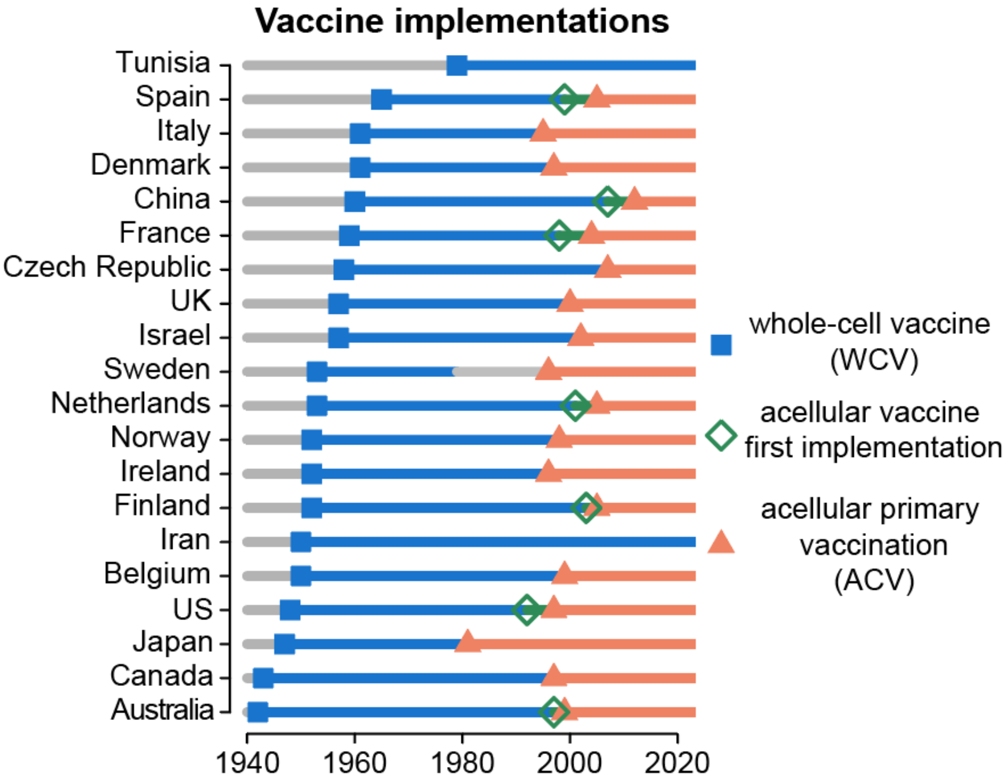

**Figure S9: Model fits for all countries.**

Fits of the proportion of each genotype for 16 countries: Belgium **(A)**, Canada **(B)**, China **(C)**, Czech Republic **(D)**, Denmark **(E)**, Spain **(F**), Finland **(G)**, Ireland **(H)**, Israel **(I)**, Italy **(J)**, Netherlands **(K)**, Norway **(L),** Sweden **(M)**, UK **(N)**, Iran **(O)**, and Tunisia **(P)**. Australia, France, USA and Japan are presented in Figure 3A-D. Data is shown in grey. Blue lines and shaded areas represent the median and 95% credible interval of the posterior. Vertical dotted line denotes the year of the first ACV introduction, for each country.

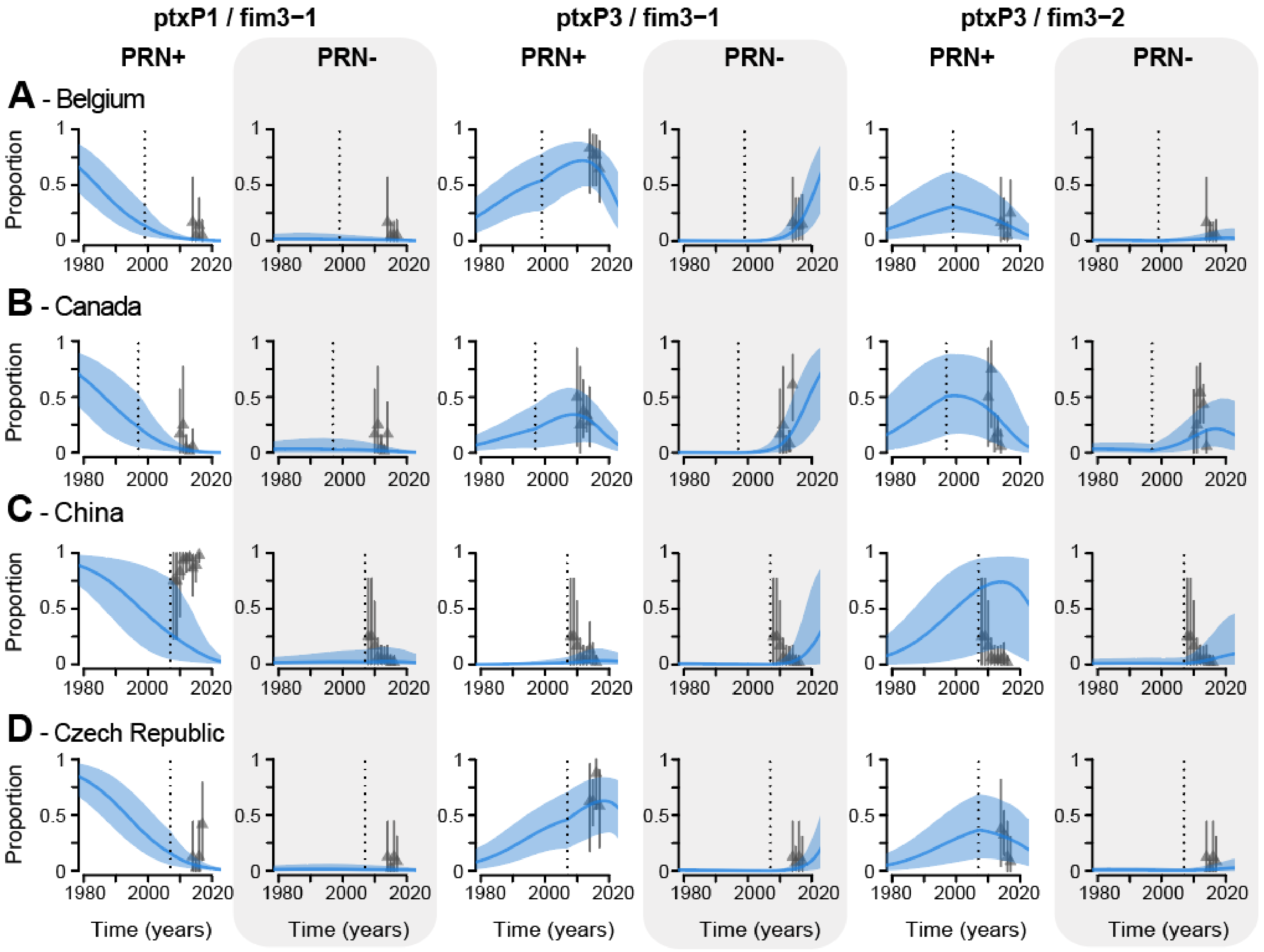

ptxP1 / fim3−1
ptxP3 / fim3−1
ptxP3 / fim3−2
PRN+
PRN-
A - Belgium
B - Canada
C - China
D - Czech Republic
Proportion
Time (years)

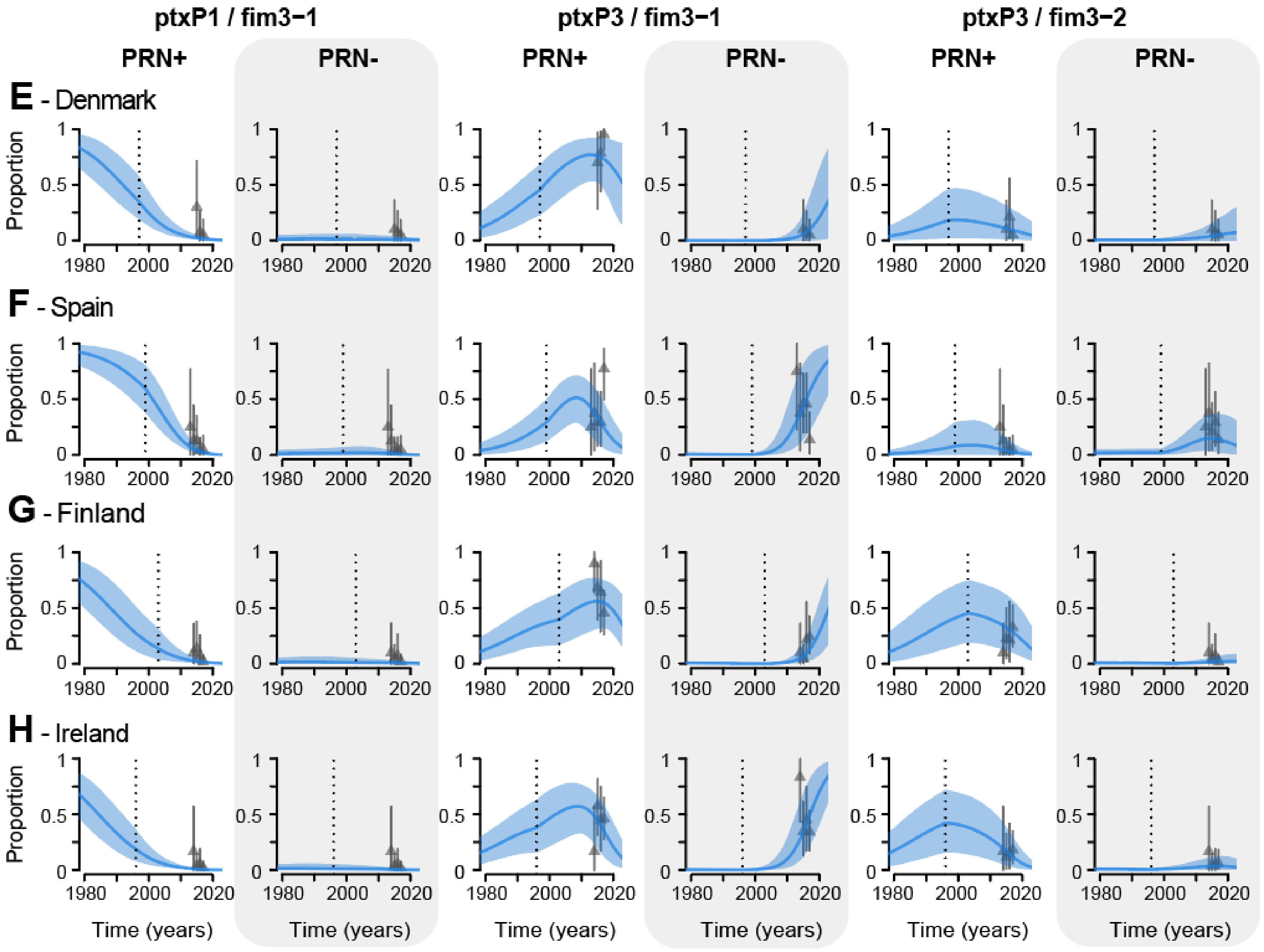
ptxP1 / fim3−1
ptxP3 / fim3−1
ptxP3 / fim3−2
PRN+
PRN-
E - Denmark
F - Spain
G - Finland
H - Ireland
Proportion
Time (years)

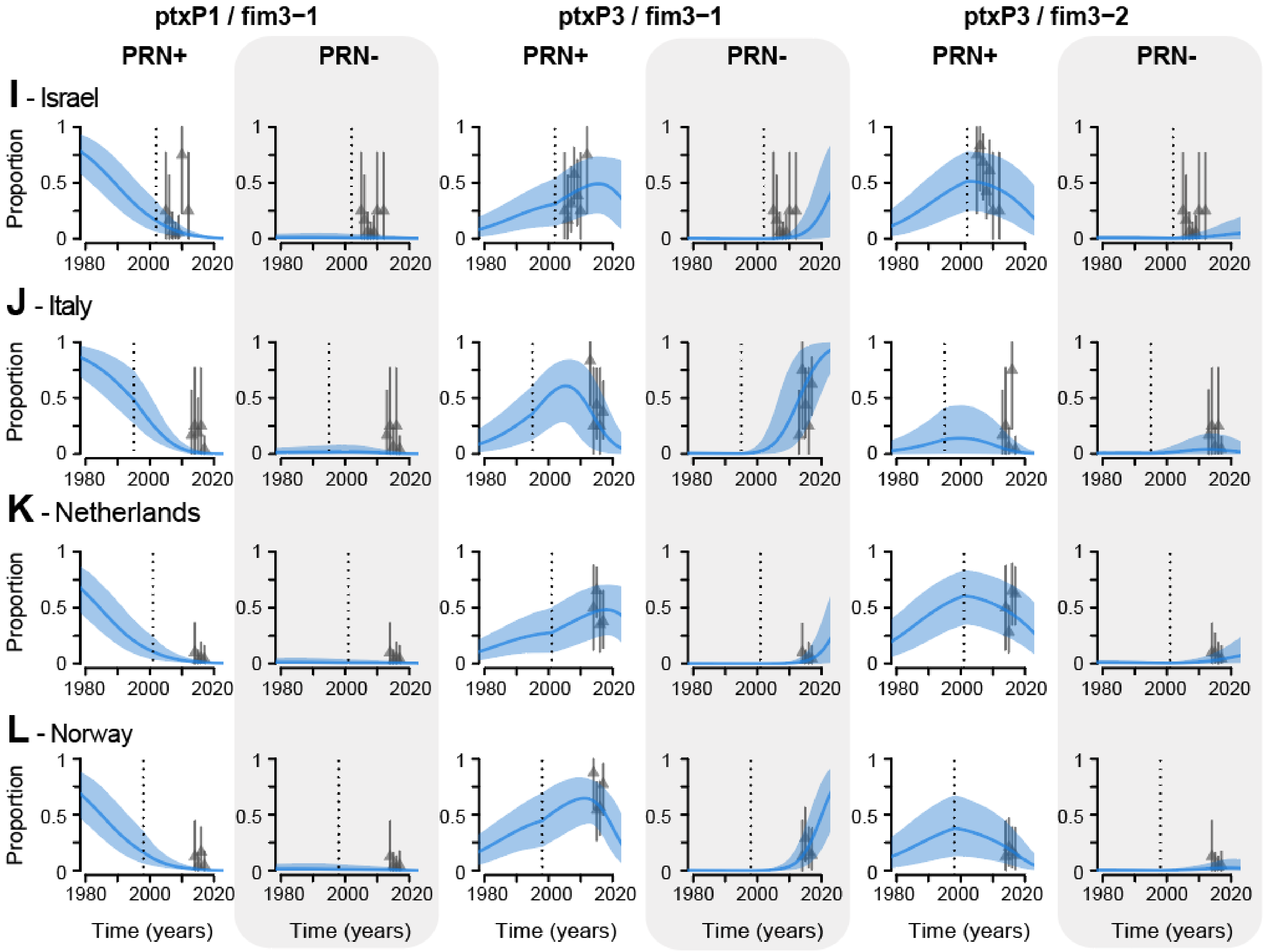
ptxP1 / fim3−1
ptxP3 / fim3−1
ptxP3 / fim3−2
PRN+
PRN-
PRN+
PRN-
PRN+
PRN-
I - Israel
J - Italy
K - Netherlands
L - Norway
Proportion
Time (years)

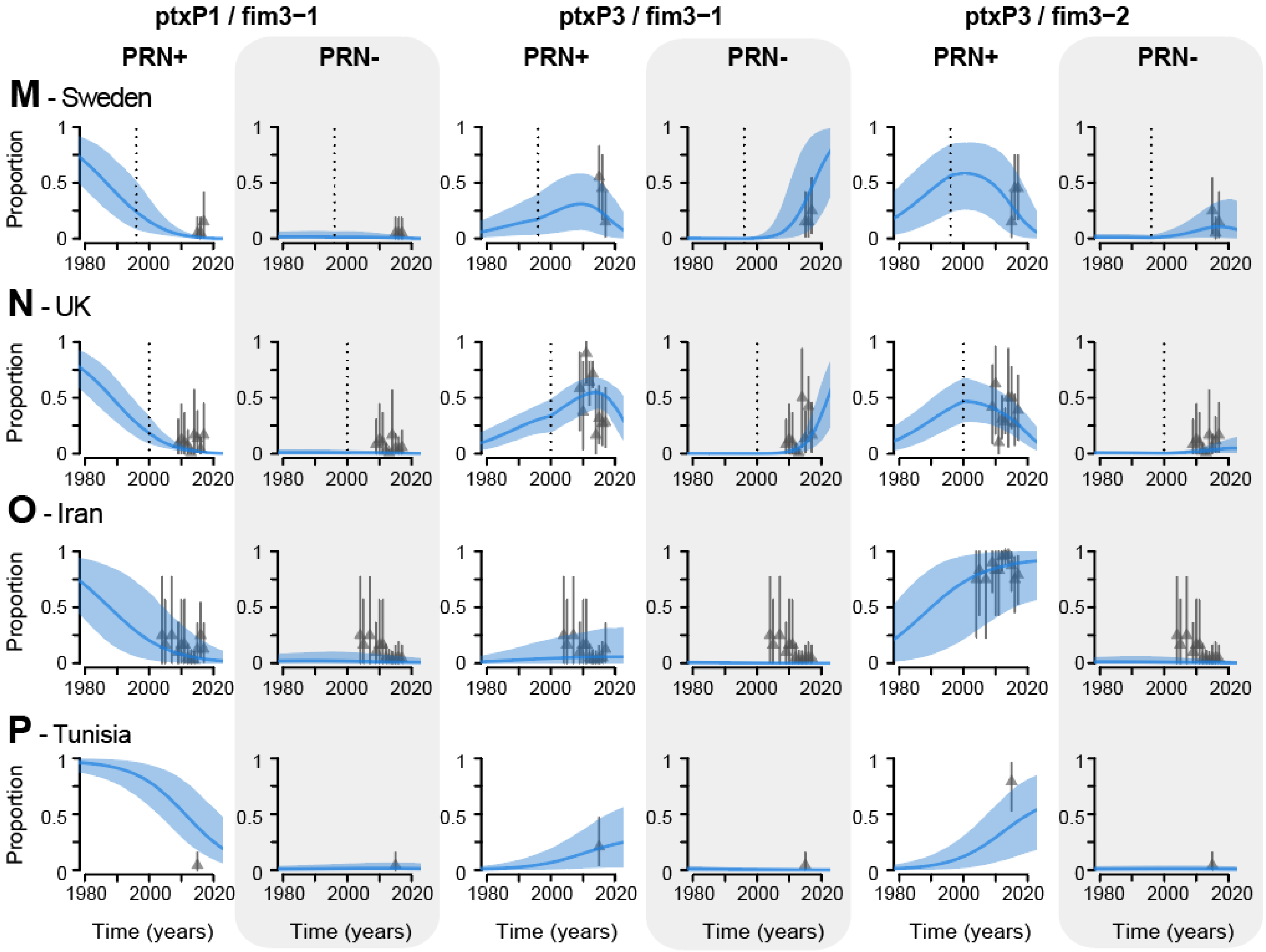
ptxP1 / fim3−1
ptxP3 / fim3−1
ptxP3 / fim3−2
PRN+
PRN-
M - Sweden
N - UK
O - Iran
P - Tunisia
Proportion
Time (years)
0
0.5
1
1980
2000
2020

**Figure S10: Model fit for different delays between ACV implementation and fitness change.**

Model adequacy has been measured with the Watanabe–Akaike information criterion (WAIC)[31]. We plot the difference to the best WAIC (y-axis) against the year of the switch in fitness (x-axis). Dashed vertical line denotes the model without any delay. The dark grey box highlights equivalent models ($\Delta WAIC <= 2$) and light grey box highlights similar models ($\Delta WAIC <= 7$) . The values for the model comparisons are presented in Table S4.

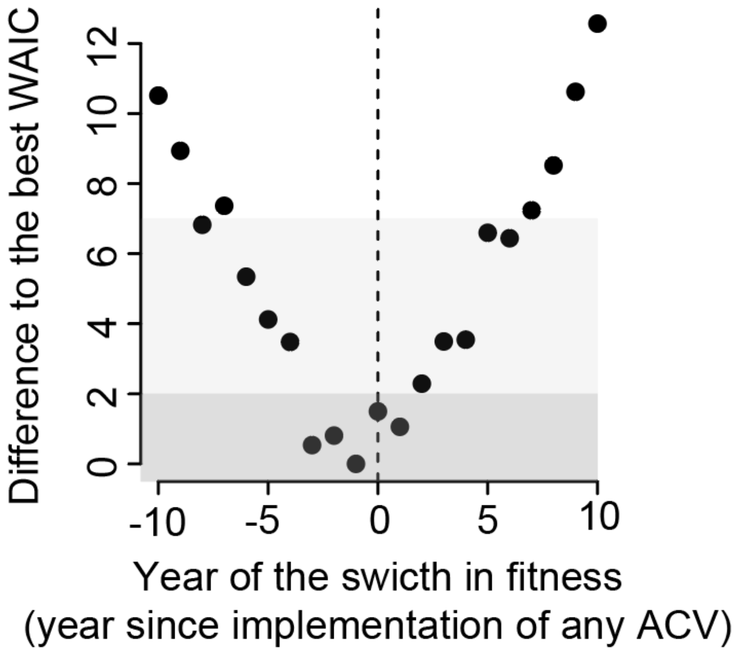

**Figure S11: Results of the simulation study for our fitness model, with or without biased sampling.**

**(A)** Proportion of cases from each strain, as a function of time, in the simulated epidemic (see supplementary materials for further details). **(B)** Sampling procedures used: for the uniform sampling the dashed line represents the number of sequences sampled each year, for the biased sampling the bar plot represents the amount of sequences sampled each year. **(C-D)** Fits of the proportion of strain 1 in the population. **(E)** Estimates of relative fitness of strain 1 versus strain 2 using either the uniform sampling or the biased sampling of the data. The biased sampling is meant to closely mimic the temporal sampling structure of our study. The green dashed line represents the true relative fitness. Dots and lines represent the median and 95% posterior credible intervals, respectively.

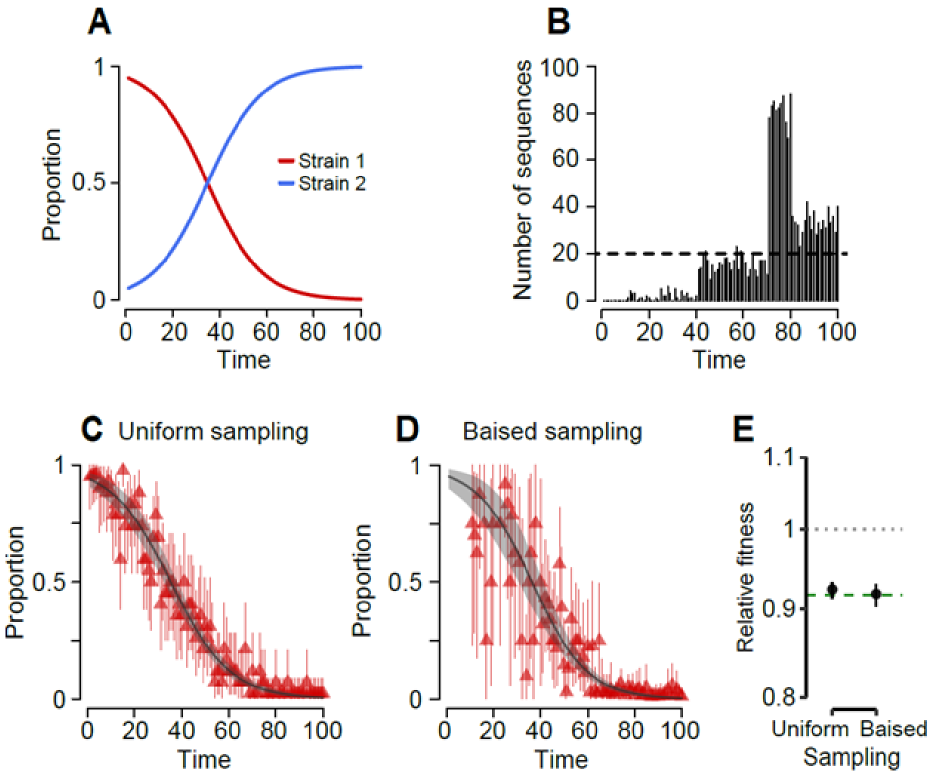

**Figure S12: Temporal signal in the dataset.**

Linear regression of root-to-tip distance (y-axis) against sampling dates (x-axis).

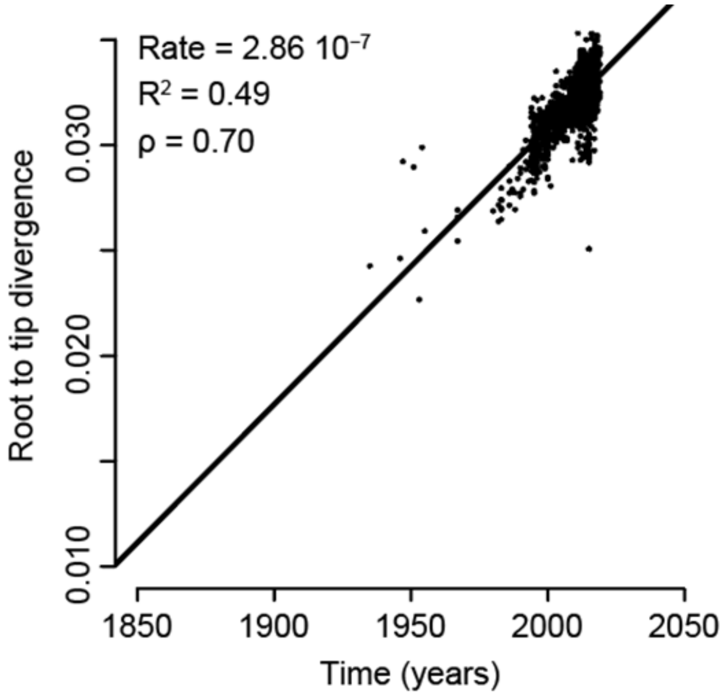

**Figure S13: Maximum clade credibility tree for all the isolates.**

Branch tips are colored by the continent of collection. The outer circle denotes pertactin (PRN) expression (black: deficient expression, grey: wild type expression, white: not known).

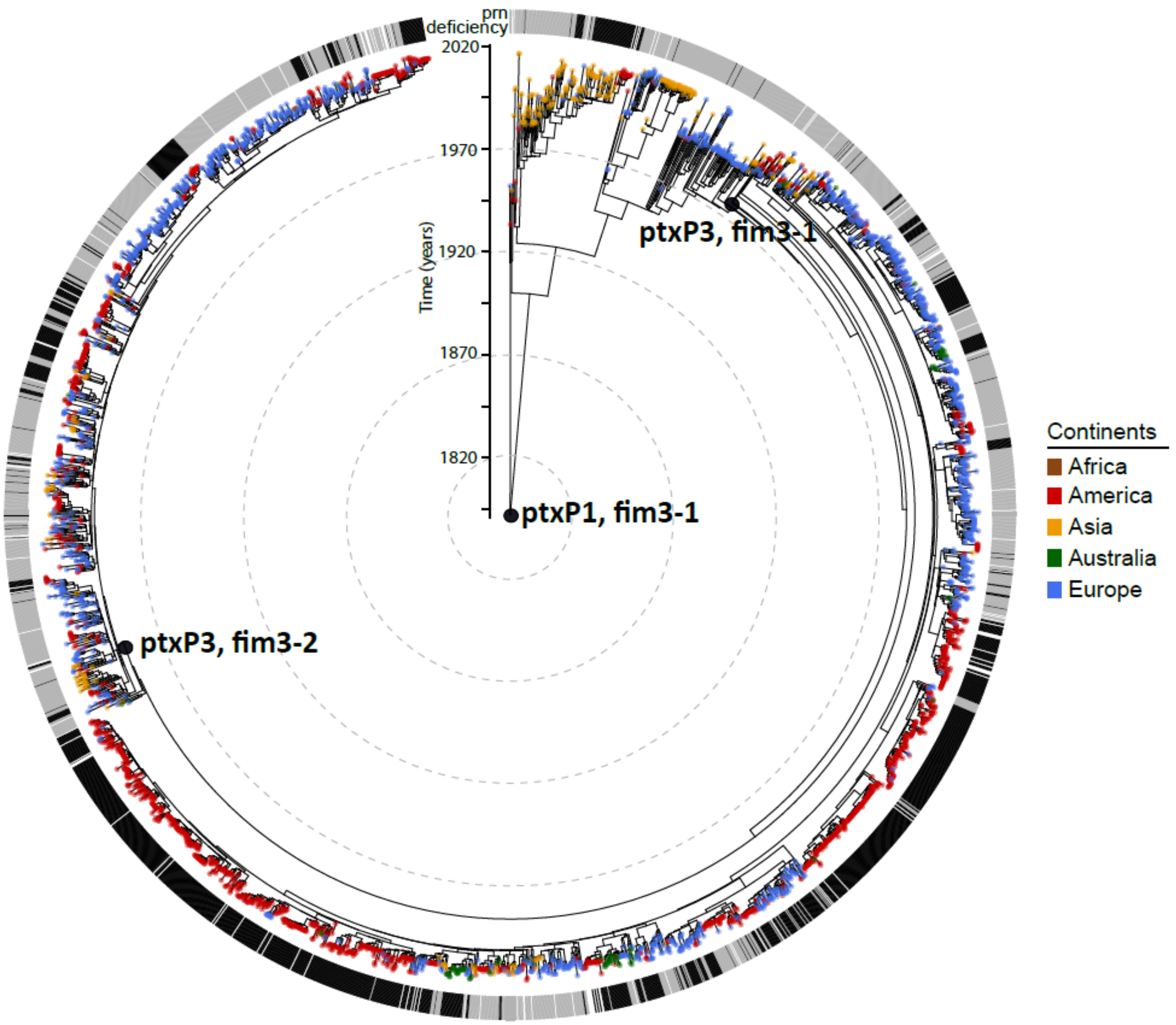

**Figure S14: Sensitivity of transmission chains estimates to changing MRCA cutoff.**
**(A and C)** Proportion of pairs within a region that belong to the same transmission chain, defined as MRCA<1y (A) or defined as MRCA<3y (C), as a function of population size (average from 19 countries). Dashed line represents model fit assuming an exponential relationship between the two and the grey shaded region 95% confidence intervals. **(B and D)** Number of transmission chains, defined as MRCA<1y (B) or defined as MRCA<3y (D), within regions, as a function of the population size. Dashed line represents model fit and the grey shaded region 95% confidence intervals.

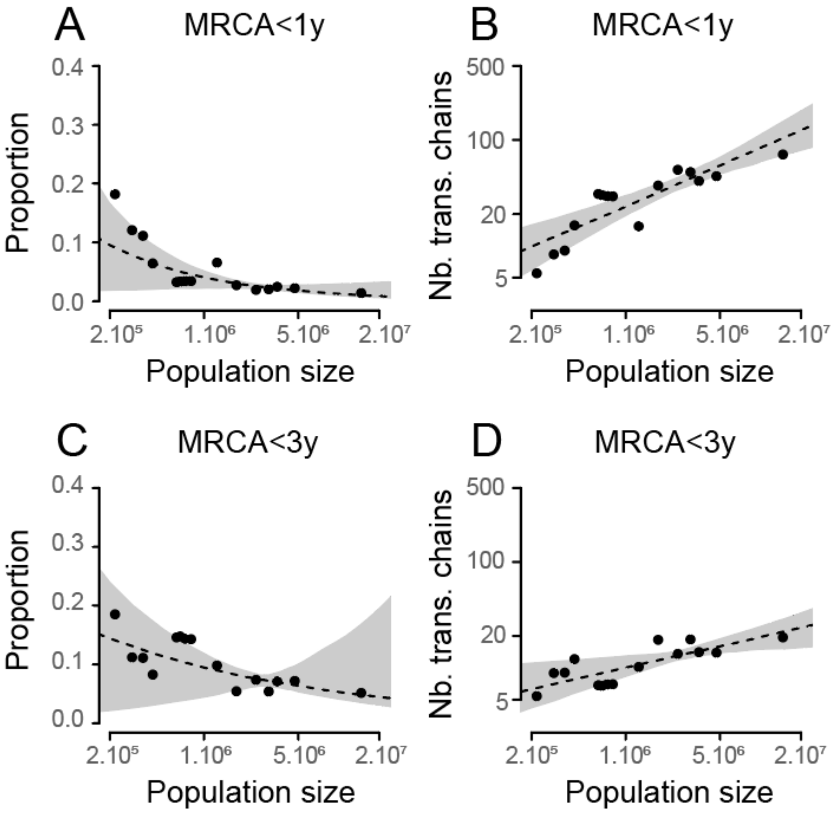

**Figure S15: Sensitivity analysis: estimates with a range of different models**

**(A)** Estimated fitness of each strain in the Whole-Cell Vaccine era (WCV), for different models tested. **(B)** Same as (A) but for the Acellular vaccine era (ACV). Dots and lines represent the median and 95% posterior credible intervals, respectively. Further details on the models tested can be found in the supplementary materials. Model comparison is presented in Table S3.

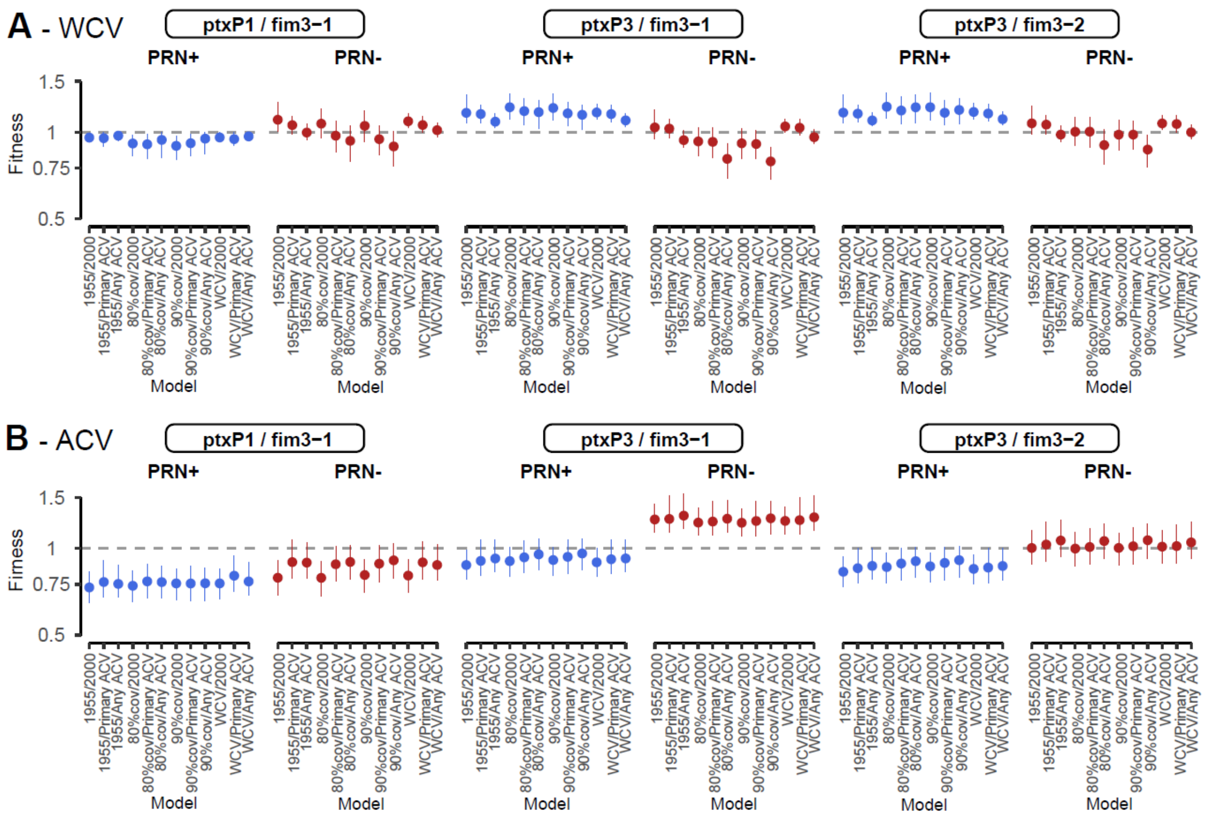

**Figure S16: Held out fitness model.**

Predicted *versus* held-out counts **(A)**, proportions **(B)**, and Wrightian fitness **(C)**, respectively. For the held-out data, 10% of the country-year data was removed from the model fitting process.

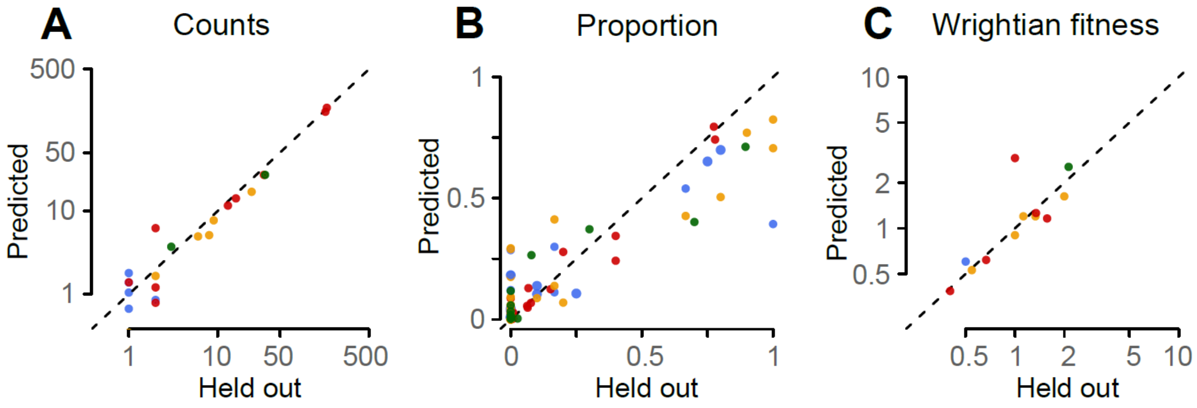
